# Growth-dependent gene expression variation influences the strength of codon usage biases

**DOI:** 10.1101/2023.03.14.532645

**Authors:** Mackenzie M. Johnson, Adam J. Hockenberry, Matthew J. McGuffie, Luiz Carlos Vieira, Claus O. Wilke

## Abstract

The most highly expressed genes in microbial genomes tend to use a limited set of synonymous codons, often referred to as “preferred codons.” The existence of preferred codons is commonly attributed to selection pressures on various aspects of protein translation including accuracy and/or speed. However, gene expression is condition-dependent and even within single-celled organisms transcript and protein abundances can vary depending on a variety of environmental and other factors. Here, we show that growth rate-dependent expression variation is an important constraint that significantly influences the evolution of gene sequences. Using large-scale transcriptomic and proteomic data sets in *Escherichia coli* and *Saccharomyces cerevisiae*, we confirm that codon usage biases are strongly associated with gene expression but highlight that this relationship is most pronounced when gene expression measurements are taken during rapid growth conditions. Specifically, genes whose relative expression *increases* during periods of rapid growth have stronger codon usage biases than comparably expressed genes whose expression *decreases* during rapid growth conditions. These findings highlight that gene expression measured in any particular condition tells only part of the story regarding the forces shaping the evolution of microbial gene sequences. More generally, our results imply that microbial physiology during rapid growth is critical for explaining long-term translational constraints.

## Introduction

In prokaryotic and eukaryotic genomes, it has long been observed that alternate synonymous codons are used non-randomly in a species-specific manner (Ikemura 1985; Plotkin and Kudla 2011; López et al. 2020). The preferential use of specific synonymous codons, also known as codon usage bias (CUB), is notably strong in highly expressed genes (Bennetzen and Hall 1982; Gouy and Gautier 1982; Sharp 1991). Strong codon usage bias is often discussed alongside observations of slow evolutionary rates in highly expressed genes, indicating that such genes may be subject to strong purifying selection (Duret and Mouchiroud 2000; Pál et al. 2001; Rocha and Danchin 2004; Subramanian and Kumar 2004). Numerous mechanistic explanations have been put forth to explain variation in codon usage biases and the correlation with expression—with causes variably attributed to mutation, selection, or drift (Hershberg and Petrov 2008; Plotkin and Kudla 2011). While the details remain unclear, there has been consistent, compelling support for translational selection as a driving force shaping the evolution of coding sequences (Sharp and Li 1987; Reis et al. 2004; Drummond et al. 2005; Drummond and Wilke 2008; Tuller et al. 2010; Park et al. 2013; Zhou et al. 2016; Hanson and Coller 2018; Frumkin et al. 2018; de Oliveira et al. 2021). Simply, highly expressed genes are subject to particularly strong selective pressures to ensure the accurate and/or efficient production of proteins, and thus have stronger bias for the codons that will maximize accuracy and/or efficiency.

The observation that gene expression is predictive of codon usage biases has been made in various species and environments but remains most pronounced in (fast growing) single-celled organisms (Pál et al. 2001; Sharp 1991; Duret and Mouchiroud 2000; Subramanian and Kumar 2004; Galtier et al. 2018; Kames et al. 2020). Most existing studies were performed under the assumption, generally not explicitly stated, that genes that are highly expressed in the specific environment in which they were measured for the purpose of the study are highly expressed in all relevant conditions. However, expression in microbes is known to be condition-dependent, varying based on current environmental conditions for example in *Escherichia coli* and *Saccharomyces cerevisiae* (Elowitz et al. 2002; López-Maury et al. 2008; Urchueguía et al. 2021). It is unclear how condition-dependent gene expression impacts expectations for codon usage bias. If a gene is expressed highly in certain conditions but not in others, the predicted strength of codon usage biases is unknown. Further, microbial populations are subjected to fluctuating environmental conditions across time and thus different selection pressures (López-Maury et al. 2008). It is unclear which conditions are of evolutionary significance in shaping gene expression levels, codon usage biases, and the relationship between them.

Here, we explore the connections between environmental condition, gene expression, and codon usage biases. We primarily focus our analysis on published transcriptome-level data coupled with known growth rates in *E. coli*, but confirm our findings in *E. coli* proteome and *S. cerevisiae* transcriptome and proteome data sets (Sastry et al. 2019; Schmidt et al. 2016; Yu et al. 2021). We find a predictive relationship between expression and CUB with the strength of this correlation varying across conditions. Conditions with the strongest correlation are consistently associated with the fastest growth rates. We show that the degree to which the expression of individual genes is correlated to growth rate—which we term the Growth Correlation Index (GCI)—is a strong predictor of CUB. We further investigate individual genes and find that those who are members of the core or essential genome are more likely to see relative expression increases during rapid growth (GCI *>* 0), while accessory and non-essential genes most often see a relative decrease in expression during rapid growth (GCI *<* 0). Finally, we complete a gene ontology (GO) term enrichment analysis for genes with positive and negative GCI values and find that resource uptake becomes relatively more important in slow growth while biosynthesis becomes relatively more important in fast growth environments. In aggregate, our findings support the notion that selective pressures on translation drive the evolution of strong codon usage biases, and that these pressures are most pronounced under rapid growth.

## Results

We explored condition-dependent gene expression by incorporating this variation into the well-studied expression–CUB paradigm. Most prior analyses examining the relationship between gene expression and codon usage bias ignored environmental variation, assuming that expression levels and the resulting dynamics with CUB are constant across all conditions. We considered a number of scenarios where gene expression profiles either conform to or violate this assumption and summarized the known expectations for codon usage patterns in these genes (Figure 1). Genes with constant expression profiles (genes A, B, and C in Figure 1) have expected bias levels that correlate with their relative expression; i.e., highly expressed genes have strong CUB, while lowly expressed genes have weak CUB. For genes with variable expression profiles (genes D and E), it is unclear what level of bias should be expected. In reality, most genes likely fall somewhere between the two extremes presented. Therefore, we characterized condition-dependent gene expression and its relationship to codon usage biases in an attempt to understand these currently unknown dynamics.

**Figure 1.**
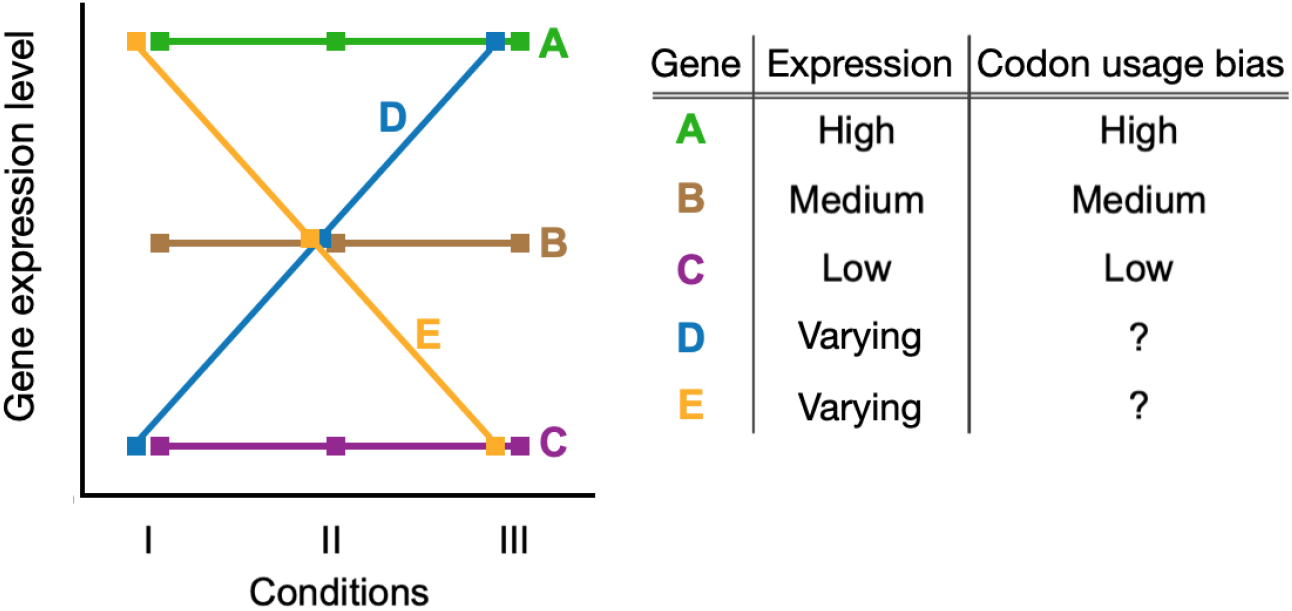
Known and unknown relationship dynamics between gene expression, growth, and expectations for CUB. Highly expressed genes tend to have strong codon usage bias (gene A), lowly expressed genes have low bias (gene C), and intermediate expression have intermediate bias (gene B). These expectations are based on observations in constant conditions and do not account for environmental variation. It is unclear what level of bias to expect for genes that have variable expression across conditions/environments (genes D and E).

### Gene expression in rapid growth conditions best predicts codon usage bias

We began our investigation into the relationship between gene expression and codon usage biases by first exploring an existing RNA-seq compendium that consists of 278 uniformly processed RNA-seq experiments performed across 154 conditions for *E. coli* (strains K-12 MG1655 and BW25113) (Sastry et al. 2019). We limited this data to conditions with known growth rates, and we averaged the expression of genes across biological replicates when applicable (see Materials and Methods). This resulted in 103 unique expression profiles whose correlations with one another (reported here as coefficient of determination, *R*^2^) ranged from 0.42 to 0.985 (Figure 2). The paired conditions with the highest and lowest reported correlation were `ytf_delydcI_ph5` vs `ytf wt_ph5 and ica_cytd_rib` vs `ssw_glc_xyl_glc`, respectively (Figure 2B). Despite the limited size of this data set and the fact that it contains only laboratory growth conditions, there was nevertheless clear and substantial heterogeneity in gene expression whose impact we sought to explore.

**Figure 2.**
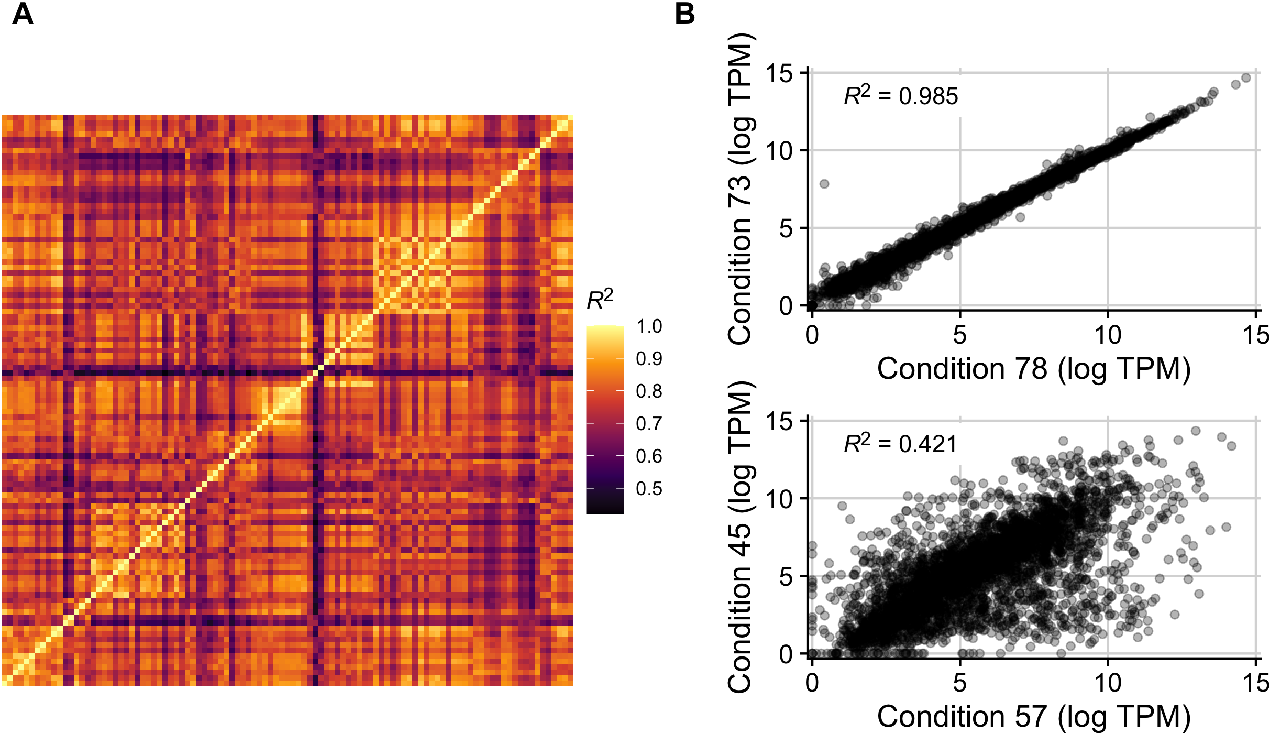
Gene expression variably correlates between conditions in the *E. coli* RNA data set with 103 conditions. (A) Coefficients of determination (*R*^2^) in an allby-all comparison of gene expression levels across all 103 conditions. Each row/column is one condition. (B) The two pairs of conditions with the highest and lowest *R*^2^ in the data set. Conditions 78 and 73 correspond to ytf_delydcI_ph5 and ytf_wt_ph5, which vary in the MG1655 strain used (one with a deletion of ydcI vs wildtype). Conditions 57 and 45 correspond to `ica_cytd_rib` and `ssw_glc_xyl_glc`, which vary in their carbon source, the presence of a cytidine supplement, and their ALE status.

We next quantified codon usage biases for all genes in each condition in the data set. We used the Codon Adaptation Index (CAI) as a measure of codon usage biases but note that results were similar when using two additional codon usage bias metrics that—including the CAI—spanned a range of assumptions: the tRNA Adaptation Index (tAI) and ROC SEMPPR (Sharp and Li 1987; Reis et al. 2004; Gilchrist et al. 2015). Across the 103 conditions in our processed data set, we observed that the strength of the relationship between CAI and measured transcript abundances varied substantially, with *R*^2^ values ranging from 0.15 to 0.30 (Figure 3A,B). The conditions with the weakest and strongest relationship between CAI and gene expression were `rpoB_rpoBE672K_lb` and `ica_cytd_rib`, respectively.

**Figure 3.**
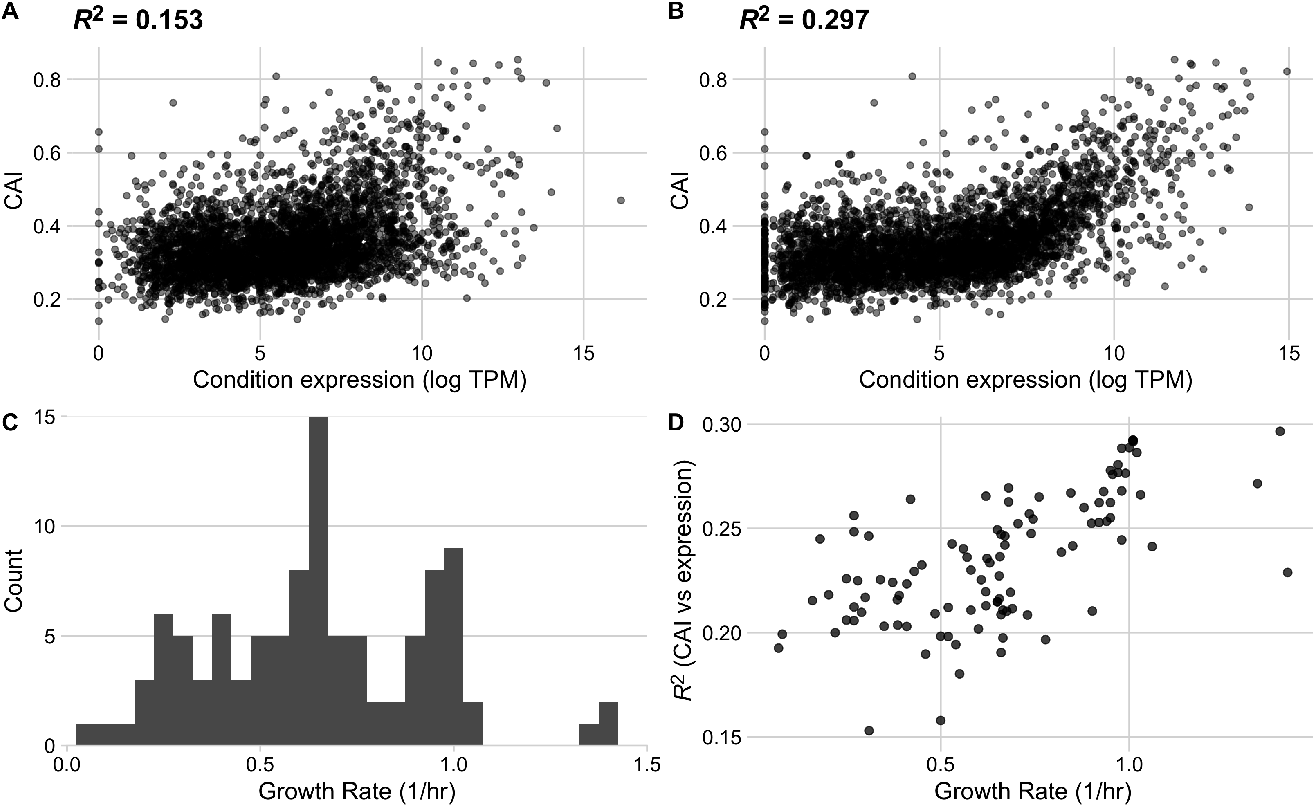
The strength of the relationship between codon usage bias and gene expression is growth-rate dependent in the full *E. coli* RNA data set. Conditions with the lowest (A) and highest (B) correlation between expression and CAI. The lowest correlation was found in condition `rpoB_rpoBE672K_lb`, a rpoB knock-in study with LB base media, a glucose carbon source and Kanamycin antibiotic, while the highest correlation was found in condition `ica_cytd_rib`, a wildtype study with M9 base media, a D-ribose carbon source, and a cytidine supplement. (C) Distribution of reported growth rates for all conditions in the assessed data set. (D) Relationship between a condition’s growth rate and the strength of the relationship between expression and CAI (reported as *R*^2^).

For quantifiable comparisons across conditions, we characterize each condition by its reported growth rate. Reported growth rates across the individual conditions vary from 0.07 to 1.42 (1/hr), corresponding to population doubling times between 0.48 and 9.9 hours (Figure 3C). We mapped the strength of the relationship between gene expression and CAI to the growth rate data and observed a strong (and significant) positive relationship: transcript abundances from rapid growth conditions were much more strongly predictive of CAI values than transcript abundances measured during periods of slow growth (Spearman’s *ρ* = 0.64, *p* = 3.30 *×* 10^*−*13^, Figure 3D).

### Individual genes vary in the extent to which they are expressed during rapid growth

We assessed the correlation between relative gene expression values across all conditions for a given gene and the corresponding observed growth rates for those conditions. The resulting correlation coefficients (Pearson’s *r*) spanned a broad range, from *−*0.77 to 0.72 (Figure 4A, B). We compared the distribution of correlation coefficients to the expected distribution under the null hypothesis of no association, by randomly permuting the data, and found that the bulk of the observed distribution fell outside the null expectation, with the largest and smallest observed correlation coefficients falling way beyond two standard deviations of the null distribution (Figure 4C). Further, the observed distribution of correlation coefficients was substantially skewed, with a mode around *−*0.3. In other words, we found that a majority of genes *decreased* in relative abundance during periods of rapid growth whereas comparatively few genes (examples include ribosomal proteins) increased in relative abundance. For ease of nomenclature, we defined the *Growth Correlation Index* (GCI) of a gene as the correlation coefficient *r* between the gene’s expression level across all conditions and the corresponding growth rates of those conditions. GCI values can span a theoretical range from *−*1 to 1, and the end points of this range represent genes whose expression is, respectively, perfectly negatively or positively correlated with measured growth rates. We found that GCI values for individual genes were positively correlated with the mean expression level across all conditions (*R*^2^ = 0.14, *p <* 10^*−*10^), indicating that the genes which tended to increase in abundance under rapid growth conditions also appeared to have higher levels of expression across the 103 conditions overall.

**Figure 4.**
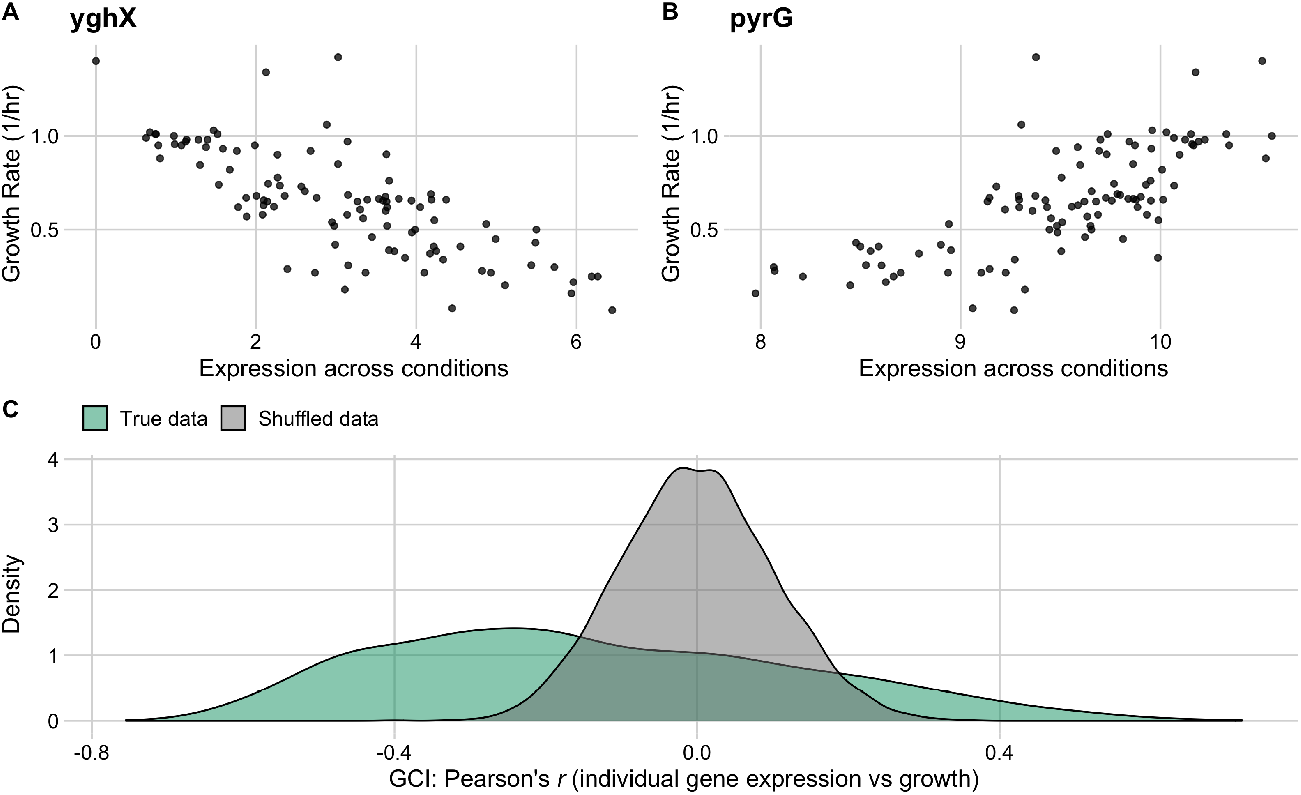
Individual gene expression across conditions variably correlates with growth rate in the full *E. coli* RNA data set. The top row shows the two genes with the most negative (A) and most positive (B) correlation between growth rate and expression across all conditions. (C) The distribution of correlation coefficients for all genes (shown in green) against a reference data set with permuted expression and growth data (grey).

### Growth-rate-dependent expression variation is predictive of codon usage bias

To test whether the GCI metric captures biological constraints, we assessed its relationship with codon usage biases. We constructed a series of linear models to predict CAI values from average gene expression or GCI values both independently and in combination (Figure 5). First, we found that the relationship between mean gene expression and CAI was highly significant and positive, as expected (adjusted *R*^2^ = 0.27, *p <* 10^*−*10^, Figure 5A, B). Second, we found that the relationship between GCI and CAI was significant and positive but slightly weaker (adjusted *R*^2^ = 0.19, *p <* 10^*−*10^). Importantly, GCI correlated more strongly with CAI than it did with mean expression level (*R*^2^ = 0.14), as reported in the preceding subsection. Also, we note that while the relationship between CAI values and mean expression for genes appeared somewhat non-linear (Figure 5B), the relationship between CAI and GCI was best characterized by a linear model (Supplemental Figure S1).

**Figure 5.**
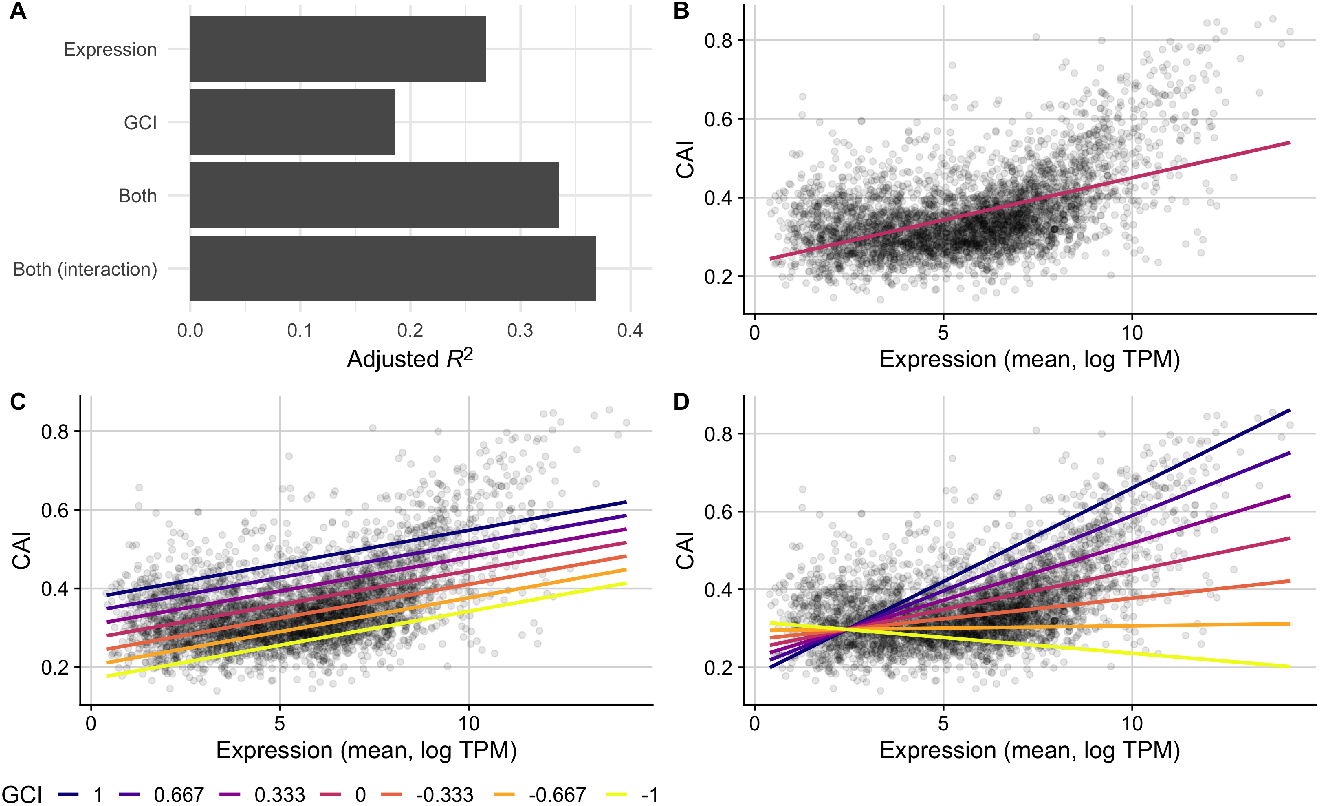
CAI is partially predicted by GCI in the full *E. coli* RNA data set. (A) A comparison of the predictive ability (measured as *R*^2^ after adjustments) of linear models that use either: 1) mean expression values, 2) GCI values, 3) both expression and GCI values, or 4) expression and GCI values with an interaction term, to predict CAI. (B, C, D) CAI against mean expression for a subset of models with observed values for each gene shown as points and model predictions as lines. The fit of model 1, which predicts CAI using only mean gene expression values, is shown with one line (B), while models 3 and 4 are shown with several lines colored by fixed GCI values (C and D, respectively).

Next, we constructed an additive multi-variable model using both GCI and mean expression to predict CAI scores. We observed a higher predictive power (adjusted *R*^2^ = 0.33, *p <* 10^*−*10^) than for either individual model, and confirmed that this increase does not appear to be caused by collinearity (VIF = 1.16). Further, we found that the overall magnitude of the contribution made by the GCI predictor variable to this model was substantial (Figure 5A, C). In practice, this shows that genes with similar mean expression levels across all conditions are likely to have substantially higher CAI scores if their relationship with growth rate is positive (GCI *>* 0). By contrast, genes that are negatively associated with growth rate (GCI *<* 0) will—on average—have substantially lower CAI scores than their mean expression across the conditions might otherwise predict. These relationships are most easily seen by comparing model predictions under different GCI values (Figure 5C).

To further ensure that the relationship between GCI and codon usage biases is robust to various model assumptions and specifications, we constructed a final model which included an interaction term between GCI and mean expression. For this model, we observed an even stronger ability to predict CAI values (adjusted *R*^2^ = 0.37, *p <* 10^*−*10^, Figure 5A, D), though the increase in variance explained was small (four percentage points). This finding shows that GCI and mean expression level are largely independent of each other in their effect on CAI, consistent with the observation of a low VIF for the model without interaction term. On average, a more positive GCI will correspond to larger CAI at all expression levels, and similarly higher expression levels will correspond to larger CAI for all GCI values.

Finally, to assess whether our results were robust to our choice of CUB metric, we repeated these analyses with tAI and ROC SEMPPR values in place of CAI. We found that regardless of CUB metric, GCI was a strong predictor of codon usage bias (Supplemental Figures S2 and S3). Moreover, for these two additional metrics of CUB, the interaction term in the fourth model had even less predictive power than it did for CAI. In aggregate, these results provide strong support that GCI is quantifying an important and previously unknown aspect of individual genes, which partially governs coding sequence evolution.

### Genes with highest GCI values are associated with important functional pathways

To gain further insight into the biological significance of GCI, we investigated what if any associations existed between GCI and the functional classification of genes. First, we checked for differences in GCI values between essential and non-essential genes. We classified all genes as either essential or non-essential based on their presence in the PEC (Profiling of *E. coli* Chromosome) database (Hashimoto et al. 2005; Yamazaki et al. 2008). We found that there was a significant difference in the distribution of the GCI values for essential and non-essential genes. GCI values for essential genes had a mode around 0.3, a mean of 0.12, and were left skewed. GCI values for non-essential genes had a mode around *−*0.3, a mean of *−*0.15, and were right-skewed. The means were significantly different (*t*-test, *p <* 10^*−*10^). We found similar results when assessing essentiality alternatively by their presence or absence in the Keio collection (Baba et al. 2006) (mean GCI values of 0.12 and *−*0.15, respectively, *p <* 10^*−*10^). We also used an additional classification where we subdivided genes into either core or accessory, rather than essential or non-essential. Core genes included those shared across 60 strains of *E. coli* (Maddamsetti et al. 2017). We found that the core genes on average differed significantly from accessory genes in their GCI values and indeed had more positive GCI values (mean GCI of *−*0.05 vs. *−*0.20, respectively, *p <* 10^*−*10^).

We also subdivided all genes by whether their GCI was positive or negative and then ran independent Gene Ontology analyses to assess GO term enrichment within each gene set. Genes with GCI *<* 0 were enriched for GO terms indicated in processes most important for cell survival, such as transport and metabolism (Figure 6B, left). By contrast, genes with GCI *>* 0 were enriched for GO terms related to biosynthesis and ribosomal activity (Figure 6B, right). We note that the odds ratios in the enrichment analysis for genes with GCI *>* 0 were considerably larger than those in the analysis for genes with GCI *<* 0.

**Figure 6.**
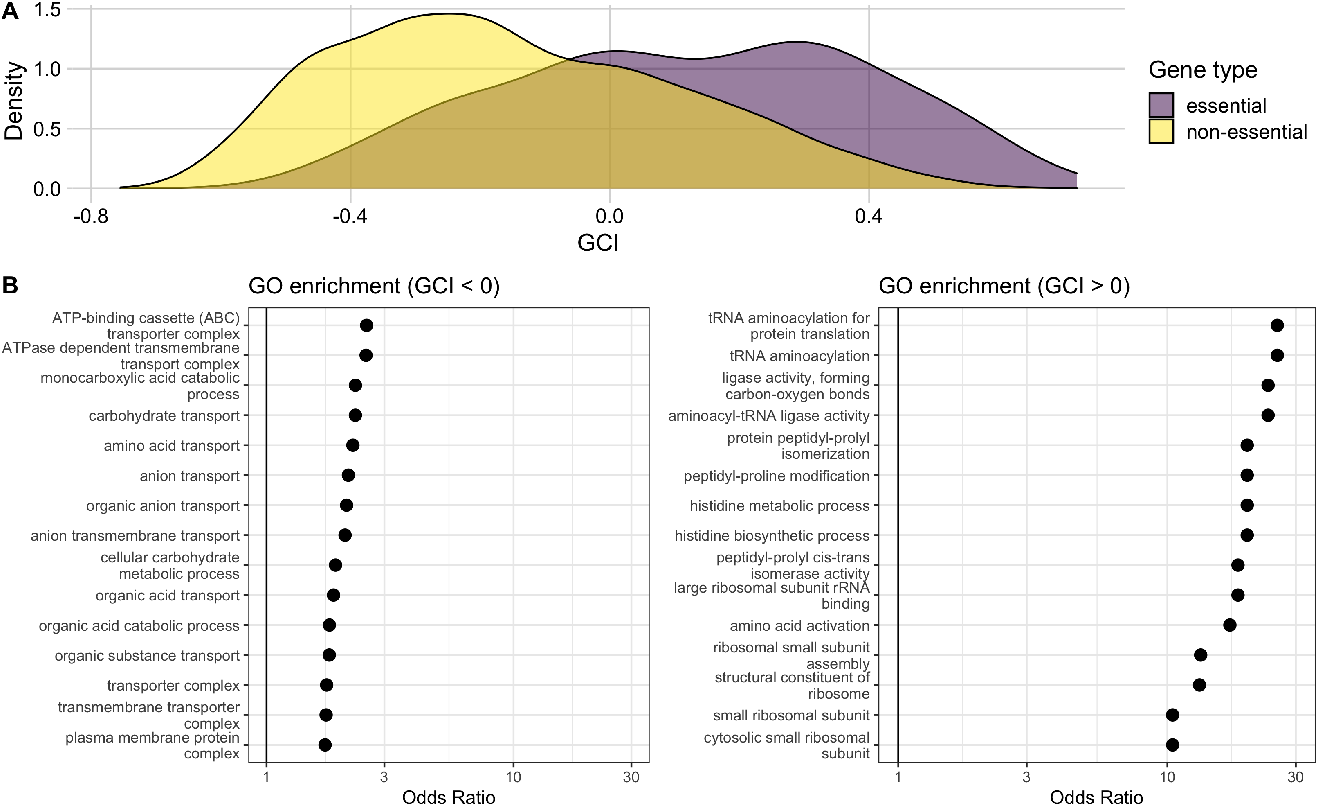
Genes with high GCI values are associated with essential genes and functional pathways. (A) Distributions of GCI values for genes classified as either essential or non-essential. The mean GCI values for these two distributions are 0.12 and *−*0.15, respectively, and they are significantly different (*t*-test, *p <* 10^*−*10^). (B) Results of a GO enrichment analysis for sets of genes with negative (left) and positive (right) GCI values. The significant terms with the 15 highest odds ratios are shown in each set. Odds ratios are shown on a log scale with a solid black reference line at 1 in each plot.

### Results are robust to changes in data sources, data filtering, and organism considered

To assess the robustness of our results, we repeated our analyses in a number of complementary data sets that covered different biomolecules (RNA and protein) and species (*E. coli* and *S. cerevisiae*). These five additional data sets consisted of two subsets of the full *E. coli* transcriptome data set, an *E. coli* proteome data set, and a transcriptome and a proteome data set from *S. cerevisiae*.

Although we had already employed several filtering steps to ensure the 103 conditions used in the full *E. coli* RNAseq data set were representative and independent, we created three subsets to account for potential limitations in the original data (Sastry et al. 2019). We constructed a “sparse” data set by iteratively removing individual conditions whose gene expression levels were strongly correlated with one another, until arriving at a set of 30 unique conditions. We additionally created two “neutral” data sets: one consisting of only the 48 conditions that were not identified as ALE (adaptive laboratory environment) strains, and the other consisting of a further subset of 28 conditions that also excluded mutants and knock-out strains. All our prior findings were qualitatively unchanged in these two data sets: GCI was highly variable, negatively-skewed, and predictive of CAI (Supplementary Figures S4, S5, S6, S7, S8, S9).

The *E. coli* genome is known to contain a subset of 730 AT-rich genes with unusually low CUB (dos Reis et al. 2003). The vast majority of these genes have negative GCI, and in fact many of the E. coli genes with the most negative GCI values are found in this group of genes (Supplementary Figure S10). Thus, these genes are among the most down-regulated under conditions of rapid growth. Therefore, by our observation that low GCI tends to associate with reduced CUB, these genes would be expected to have among the least codon usage bias in the E. coli genome. We asked whether these genes were responsible for the genome-wide relationship we have observed between GCI and CUB, by excluding them from the analysis, and found that the overall patterns remained unchanged even after excluding these genes (Supplementary Figure S11).

Next we expanded our analysis from transcript abundances to protein abundances. It is known that transcript abundances are not necessarily predictive of protein abundances *E. coli*, in particular during periods of starvation (Houser et al. 2015), so this additional analysis served as an important additional control. We utilized data that measured growth rate and protein abundances of *E. coli* across 20 conditions, where population growth rates spanned a range from 0.12–1.9 hr^*−*1^ (Schmidt et al. 2016). Despite the smaller size of the data set and the narrower range of the growth rates explored, our results were again qualitatively unchanged. Notably, protein abundances measured during conditions of rapid growth showed a stronger relationship with codon usage biases than abundances measured during slow growth. Gene-specific GCI values calculated from this data set spanned a wider range of magnitudes with a less extreme shift towards negative values (Supplementary Figure S12), and were predictors of CAI independently of expression level (Supplementary Figure S13).

We also considered *S. cerevisiae*, a well-studied microbial eukaryote. Yu et al. (2021) measured both transcript and protein abundances for thousands of *S. cerevisiae* genes across 22 conditions. We focused our analysis on a subset of those conditions encompassing 14 unique environments (see Methods). The growth rates within these conditions spanned a range of 0.05–0.35 hr^*−*1^, and correlations between individual transcriptome (proteome) data sets across all conditions spanned a range from 0.57–0.99 (0.88–0.99 in proteomes).

Qualitatively, all of the results that we observed for *E. coli* remained true for *S. cerevisiae*. Most notably, expression measurements from rapid growth conditions showed the strongest correlations with CAI values. GCI values spanned nearly the full range of potential values (*−*1 to 1) with peaks forming towards more extreme values (Supplementary Figures S14, S15). While there were only 14 unique conditions used to estimate GCI values for individual genes, the relationship between GCI and CAI was highly significant and independent of the relationship between mean expression and CAI (Supplementary Figures S16, S17).

## Discussion

We have investigated the relationship between condition-dependent gene expression and codon usage bias (CUB). We have confirmed the well known observation that CUB increases on average with increasing expression level, and we have additionally discovered that CUB is best predicted by expression values measured under conditions of rapid growth. The expression level of individual genes varies considerably across conditions and can be variably associated with microbial growth rate. To capture this association between variation in gene expression and variation in growth rate across conditions, we have introduced a novel gene-level metric, GCI (Growth Correlation Index), and we have found that GCI has a significant, positive relationship with CUB. We have further found that these results are consistent across different data sets, covering different unicellular organisms (*E. coli* and *S. cerevisiae*), different types of biomolecules (mRNA transcripts and proteins) and different metrics of codon usage bias (CAI, tAI, ROC SEMPPR). In all cases, genes with positive GCI values experience a relative increase in expression during periods of rapid growth (by definition) and they have the strongest codon usage biases. Additionally, we have shown for *E. coli* that these genes are more likely to be classified as essential genes or as members of the core genome with functional roles in translation.

Our discovery that an additional, expression-related metric (GCI) is predictive of the strength of codon usage biases, independently of the mean gene expression level across conditions, offers both renewed support and additional clarification for prior work noting the positive correlation between expression level and CUB (Pál et al. 2001; Sharp 1991; Duret and Mouchiroud 2000; Subramanian and Kumar 2004; Galtier et al. 2018; Kames et al. 2020). In these earlier studies, the observed correlations were significant but somewhat moderate. We have here observed a strong increase in our ability to predict CAI (and other CUB metrics) by accounting for additional sources of variation related to growth rate. Previously calculated correlations might have been considerably higher if growth-rate-dependent expression variation had been accounted for as we have done here.

We emphasize, however, that our work is based on simple correlation metrics, whereas several prior works have employed mechanistic population genetics models to disentangle the effects of mutation, selection, and drift on codon usage bias (Bulmer 1991; Shah and Gilchrist 2011; Wallace et al. 2013; Gilchrist et al. 2015). Importantly, all these models are steady-state models and they implicitly or explicitly assume the existence of a latent gene expression level that is constant over time. Consequently, there is no straightforward way to adapt them to the inherently non-equilibrium scenario we are considering here, where we have no *a priori* knowledge about how expression changes relate to growth rate and what distribution of growth conditions species experience over evolutionary time. We believe that given this lack of *a priori* knowledge, an unbiased, correlation-based approach is the right tool to discover novel biological relationships, such as the observation we have made here that an increase in expression level with increasing growth rate seems to lead to the strongest selection pressure on codon usage. We hope that future modeling studies will take this observation and incorporate it into a mechanistic model of mutation, selection, drift, and growth under changing environmental conditions.

During periods of rapid growth, genes with GO terms associated with biosynthesis pathways experience a relative increase in expression. In addition to being upregulated, these genes also display strongest correlations between the growth-dependent expression level (as quantified by GCI) and codon usage bias. The observation that this pattern holds across species, biomolecules, and metrics of codon usage bias provides strong support for the notion that highly expressed genes in high growth environments experience the strongest selection. While there could be many biological explanations, we suggest one that we believe to be most likely: Under conditions of rapid growth, microbial cells risk depleting their charged tRNA molecules, and the genes that are highly expressed under these conditions (genes associated with biosynthesis and essential functions) experience the strongest selection to preferentially use the tRNAs (and thus codons) that will be the least limiting to growth. This explanation aligns with insights from previous modeling studies (Sharp and Li 1987; Reis et al. 2004; Tuller et al. 2010; Hanson and Coller 2018; Frumkin et al. 2018; de Oliveira et al. 2021). We emphasize that this explanation does not specifically distinguish between selection for speed or accuracy of translation, nor does it provide any insight into whether selection is driven primarily by the rate of protein production, the absolute amount of protein produced, the rate or amount of misfolded protein produced, or any other biological function involved in accurate and efficient translation under rapid growth.

Our observation that GCI (an empirically determined metric quantifying the correlation between individual expression and growth rate across conditions) correlates strongly with several metrics of codon usage bias is complementary to previous observations made at the species-level (Sharp et al. 2010). Sharp *et al*. surveyed 80 bacterial species and found that species-level growth rates and codon usage biases were highly correlated. Bacterial species with stronger biases had a higher number of rRNA operons and tRNA genes in their genomes (Sharp et al. 2010). This work was confirmed and extended in numerous studies (Vieira-Silva and Rocha 2010; Ran and Higgs 2012; Hockenberry et al. 2018; Weissman et al. 2021). While species-level patterns have been well established, our analysis shows that growth rates and codon usage biases are highly correlated within individual species as well.

We have found through an enrichment analysis on GO terms that resource uptake tends to become relatively more important in slow growth environments (GCI *<* 0) whereas processes such as biosynthesis become relatively more important in fast growth environments (GCI *>* 0). The genes responsible for these processes experience a relative increase in expression in those contexts. This trend is consistent with prior work on starvation in *E. coli* (Houser et al. 2015) and with the observation that growth-related and stress-related genes have antagonistic regulation (López-Maury et al. 2008). In addition to implications in translational machinery, genes with positive GCI values are more likely to be classified as either core or essential in reference data sets (Hashimoto et al. 2005; Yamazaki et al. 2008; Baba et al. 2006; Maddamsetti et al. 2017). Core genes are defined based on conservation across species or strains found in a variety of environments (Maddamsetti et al. 2017), and essential genes are defined based on the survival (or not) of gene knockout strains. Although there is overlap between these two sets of genes they are distinct, and genes in both sets are more likely to be upregulated during periods of rapid growth (GCI *>* 0).

Despite the abundance of available transcriptome and proteome scale data sets that have been produced in recent years, few studies have explicitly grown cells in different environments while noting the population growth rates or doubling times. This is partially a logistical challenge since growth rates are often calculated from a series of points during exponential growth; it is unclear how to explicitly extend our methods to include data from growth conditions where the cells are not growing exponentially, as may be the case during nutrient up- and down-shifts or during extremely slow stationary phase growth periods. The data sets that we selected for our analysis were all comparatively recent, spanned a range of conditions, and within each data set were uniformly processed (Sastry et al. 2019; Schmidt et al. 2016; Yu et al. 2021). We focused here on two unicellular organisms: a representative prokaryote, *E. coli*, and a representative eukaryote, *S. cerevisiae*. Of course, the data sets we used are of limited size and environmental diversity, but given the large evolutionary distance between *E. coli* and *S. cerevisiae* we expect that our results will generalize to a larger set of species.

Throughout this work, our primary data source was Sastry et al. (2019), who curated a collection of 250 RNAseq data sets for *E. coli* grown in various environments. While we have filtered this data set to ensure maximum reliability of our analysis, we acknowledge that there are still potential limitations to this data. One potential concern arises around the inclusion of adaptive laboratory environment (ALE) and mutant and knock-out strains in the 103 conditions considered. Our results have proven quantitatively consistent in the presence and absence of ALE, mutant, and knock-out strains, suggesting that their inclusion does not impede the applicability of this analysis. We note that one mutant strain is highlighted as the condition with the weakest recorded relationship between CAI and gene expression (Figure 3A); however, we emphasize that this result is presented only as an example of a weak correlation.

We additionally note that gene expression and cellular growth rate measurements are inherently noisy. And some of the noise is due to measurement error while other noise represents biologically meaningful variation among related organisms or across time in response to the same environment. In prior work, using a steady-state model, Wallace et al. (2013) have been able to disentangle different noise sources from each other and from mutation and selection pressures acting on codon-usage bias. However, it is not clear how to adapt this model to a non-equilibrium setting, as we have mentioned above. Furthermore, we note that expression noise is condition-dependent, tied closely to the structure of gene regulatory networks, and has been shown to systematically decrease with growth rate at the genome-level (Urchueguía et al. 2021). The relationship between noise in expression and in growth rate appears to be the target of natural selection (Urchueguía et al. 2021; Krah and Hermsen 2021). Highly expressed genes experience minimal expression noise, and have the potential to drive growth rate when fast growth is selected for (Krah and Hermsen 2021).

It is unclear what impact, if any, noise might have on our analysis, but we would expect that conditions of higher expression noise lead to weaker correlations between expression level and codon usage bias, both because the noise weakens the ability of selection to act on codon usage bias and because it prevents us from obtaining precise measurements of gene expression level. The quantity we use to study the relationship between expression level and growth rate, GCI, is a correlation coefficient between two noisy quantities, and thus is expected to be noisy as well. Nevertheless, for individual genes the GCI index should roughly capture the component of gene expression variation that is attributable to growth rate variation, as long as a reasonable number of conditions are sampled. Inaccuracies in measuring GCI are, if anything, expected to dilute statistical signal. Consequently, our findings likely represent a lower-bound regarding the strength of the relationship between growth-dependent gene expression variation and codon usage biases.

Finally, we would like to emphasize that measures of codon usage bias such as CAI, tAI, or ROC SEMPPR assume that codon preferences are constant across different environmental conditions, but tRNA pools and codon preferences are known to vary somewhat in response to growth conditions or external stressors (Dong et al. 1996; Novoa and Ribas de Pouplana 2012; Chionh et al. 2016; Torrent et al. 2018). For example, a seminal study demonstrated that in *E. coli*, abundances of tRNAs corresponding to preferred codons increased with increasing growth rate (Dong et al. 1996). It is not surprising to see tRNAs upregulated when they are the most needed, and this dynamic could also increase the selection pressure for genes expressed under rapid growth to be encoded using those preferred codons, consistent with our findings here. However, we need to be careful to assume this is a foregone conclusion. The selection strength under specific conditions will depend on the balance between the availability of and the demand for specific tRNAs, and if increased tRNA abundances cannot compensate for increased demand then selection might actually favor less biased codon usage under such conditions. Because of these subtle complexities, we see an opportunity and need to develop models that explicitly link environmental conditions, dynamic tRNA regulation, and selection on codon bias, so that we can develop better intuition and insight into how varying growth conditions shape selection for codon usage bias.

## Materials and Methods

### Data sources and processing

We analyzed several data sets that contain transcript or protein abundance data across multiple conditions for two microbes, *E. coli* and *S. cerevisiae* (Sastry et al. 2019; *Schmidt et al. 2016; Yu et al. 2021). These include three E. coli* transcriptome data sets (a full, a sparse, and a neutral version), an *E. coli* proteome data set, and *S. cerevisiae* transcriptome and proteome data sets. Analyses presented here were run independently on each data set. The original data sources and processing steps are described below.

#### *E. coli* transcriptome data

We used a previously published *E. coli* data set that contains RNA-seq and meta data for several hundred independent experiments in strains K-12 MG1655 and BW25113 (Sastry et al. 2019). Transcript abundances in this data set were reported as log-TPM (transcripts per million) throughout, and growth rates were reported as increase per hour. We limited our analysis to the experiments with reported, non-zero growth rates; i.e., we removed conditions where growth rate data was either unknown or reported as zero. Growth rates reported as zero could either indicate that stationary phase cultures were used or that errors occurred in reporting. Additionally, we excluded conditions that were reported to have poor alignment scores (less than 80). After these quality controls, 173 of the original 278 conditions remained.

Many of the experiments included in this data set had the same replicated conditions, resulting in multiple gene expression and growth measurements for one condition. To incorporate as much data as possible while avoiding pseudoreplication, we averaged both gene expression levels and growth rates across all replicates within each condition (identified within the data set based on the naming conventions given in the original paper). After averaging across replicates, 105 conditions remained. We note that two columns within the gene expression data set were identical and appeared to be duplicated (namely, conditions `pal_lyx_ale2` and `pal lyx_ale4`). Identical gene expression measurements were identified by correlation coefficients of 1.0 in an all-by-all Spearman correlation, which is extremely unlikely to arise experimentally. Despite duplicated measurements in the expression columns, the reported growth rate for each condition varied. In this case, neither averaging nor eliminating one of the duplicate columns seemed appropriate. We thus excluded these two conditions from all analyses, resulting in a final *E. coli* gene expression data set for 3,923 genes across 103 independent conditions.

While we had implemented a number of filters to ensure that conditions were unique, we created two additional data sets where filtered even further. To create a data set with maximally distinct conditions, we pruned the full data set using an iterative removal process. In each iteration, an all-by-all Spearman correlation matrix was calculated, the two conditions with the highest correlations were identified, and then one of them was randomly removed. This process continued until a data set with only 30 conditions remained. We refer to this pruned data set as the “sparse” data set. We additionally created two “neutral” data sets. The first excluded all conditions that were identified as adaptive laboratory environment (ALE) strains, limiting the data set to 48 of the 103 conditions. The second further restricted the non-ALE neutral data set to also exclude mutants and knock-out strains, reducing our analysis to 28 conditions.

Finally, for our full *E. coli* gene expression data set, we considered the impact of genes previously identified as AT-rich outliers (dos Reis et al. (2003); Table S1, group 3 genes), by repeating our analysis on a reduced data set that removed those genes. The reduced data set included 3,339 non-outlier genes measured across all 103 conditions.

#### *E. coli* proteome data set

Our *E. coli* proteome data set was taken from a study that measured protein abundance levels across 22 experimental conditions (Schmidt et al. 2016). We focused here on the measurements collected for *E. coli* strain BW25113, including only the conditions that had non-zero growth rates recorded. We limited our analysis to proteins with measured (i.e., non-missing) abundance values across all conditions and removed any duplicated expression profiles. Our final data set consisted of 2,052 unique proteins observed across 20 conditions.

Because protein abundances were reported as average counts of protein copies per cell, we transformed them to units comparable to log TPM, using the formula 6 + ln(*a/* ∑*a*), where *a* is the abundance (raw count) for a given protein within a condition, and ∑*a* sums the abundances of all proteins within a condition.

#### *S. cerevisiae* transcriptome and proteome data sets

We analyzed transcript and protein abundance data from a study that quantified expression profiles for yeast in 22 steadystate chemostat cultures (Yu et al. 2021). These 22 cultures consisted of 14 experiments (referred to as novel conditions) that were all collected using a consistent experimental protocol and 8 experiments that had been conducted previously. We focused here exclusively on the 14 novel conditions.

This yeast data set reported both transcript and protein abundances for the same conditions. Abundances for both proteins and mRNA were reported as absolute measurements in fmol/mgDW. As in the case of the *E. coli* proteome data set, we therefore normalized all abundances using the formula 6 + log_10_(*a/* ∑*a*). Additionally, we replaced any counts of zero with the smallest, non-zero minimum value in the set prior to this transformation. For the RNA data, no replacement was necessary, while the protein data required 1 replacement of 0 with 0.0218. Each experimental condition included 3 biological replicates, which were averaged (after normalization) to give one expression profile per condition.

The reported dilution rates were assumed to be equivalent to and treated as growth rate per hour in our analysis.

### Gene metrics

To perform sequence-level analyses such as calculations of codon usage bias we used reference genomes for *E. coli* (str. K-12 substr. MG1655, accession NC 000913) and *S. cerevisiae* (str. CEN.PK113-7D assembly ASM26988v1, accession GCA 000269885.1) from GenBank and Ensembl Fungi, respectively(Sayers et al. 2021; Howe et al. 2021). We limited analyses to coding sequences within each genome. For each coding sequence, we identified the locus tag, gene name, start and stop loci, and the strand it is located on. We next applied some filtering rules to ensure all genes met basic quality standards. We only included coding sequences in the analysis if they met the following criteria: First, coding sequence length had to be divisible by 3 and equivalent to the counted base length. Second, genes with unrealistically long coding sequences, defined as longer than 5 times the median length across all genes, were excluded from analysis. This process eliminated 58 coding sequences from the *E. coli* genome and 53 coding sequences from the *S. cerevisiae* genome, resulting in reference data sets with 4,357 and 5,398 genes, respectively.

For each gene in the two reference genomes, we calculated several codon usage bias metrics: the codon adaptation index (CAI), the proportional differences in ribosome overhead costs in a Stochastic Evolutionary Model of Protein Production Rate (ROC SEMPPR), and the tRNA adaptation index (tAI) (Sharp and Li 1987; Gilchrist et al. 2015; Reis et al. 2004). CAI measures the distance of a given coding sequence from a pre-defined, species-specific reference set specifying weights for each codon. It is calculated as the geometric mean of the weights for all codons within that coding sequence. Reference weights were taken from the original paper (Sharp and Li 1987). ROC SEMPPR is a CUB metric with two unique advantages: it is grounded in population genetic theory (allowing for the contributions of natural selection and mutational biases to be disentangled) and its calculations are performed with only the set of coding sequences of interest (removing the requirements of *a priori* information) (Gilchrist et al. 2015). We implemented the ROC SEMPPR model in R using the AnaCoDa package (Landerer et al. 2018). Lastly, tAI metric is a measure introduced to test translational selection by estimating the adaptation of a gene to the genomic tRNA pool (Reis et al. 2004). Like CAI, is calculated by assigning each codon a value based on a list of codon weights and then calculating the geometric mean of all weights for a given gene. Reference codon weights for *S. cerevisiae* and *E. coli* were taken from published tables (Reis et al. 2004; Tuller et al. 2010).

### Relationship between gene expression levels and growth condition

To assess the variation in gene expression levels across conditions, we computed Pearson’s correlation coefficients (*r*) and their squares, the coefficients of determination (also written as *R*^2^), for each possible gene pair. These computations were performed on the log-TPM values for mRNA abundances and on equivalently transformed values for protein abundances (see Data sources and processing).

Additionally, for each individual gene, we calculated the Pearson correlation coefficient *r* between expression level and growth rate across all conditions. We refer to this correlation coefficient as the *Growth Correlation Index* (GCI). GCI has a potential range between *−*1 and 1. Genes with negative GCI values are upregulated during periods of slow growth while genes with positive GCI values are upregulated during periods of rapid growth.

As a control, for each data set we also generated a set of randomized GCI values. These values were generated via permutations; i.e., expression values and growth rates were shuffled. For each gene, we shuffled the measured expression values and growth rates and calculated one permuted GCI value from the randomly paired data.

The set of AT-rich genes identified by dos Reis et al. (2003) was shown to have low CUB, so we compared the distribution of GCI values for these genes against the rest of the genes in our data set. We extracted gene names from Table S1, group 3 genes of dos Reis et al. (2003) and identified 508 AT-rich genes in our full *E. coli* data set. (While the original list of AT-rich genes contains closer to 750 genes, not all of them were present in our full data set.) We tested for significant differences in GCI between these 508 genes and the remaining 3,339 genes using Student’s t-test.

### Predicting Codon Usage Bias

We tested to what extent gene expression level and/or GCI predict codon usage bias, as measured by CAI, tAI, or ROC SEMPPR, by constructing a series of linear models that had the measure of codon usage bias as the response and either gene expression level or GCI or both as predictors. Specifically, we considered four different linear models. The first only used gene expression level as predictor. The second only used GCI as predictor. The third used both gene expression level and GCI as independent, additive predictors. The fourth also included an interaction term between gene expression level and GCI. For all models, we used adjusted *R*^2^ values as a measure of model performance. To assess whether the third model (with two independent variables mean expression level and GCI) suffered from collinearity, we calculated the variance inflation factor (VIF) for this model (Fox and Weisberg 2019). VIFs near one indicate no collinearity whereas VIFs in excess of 5 are usually considered problematic. The VIFs we observed were ¡2 in all cases.

### Relationship between GCI and functional annotations of genes

For additional insight into the biological significance of our GCI metric, we examined its relationship to functional properties of genes. We performed this analysis only for the data set corresponding to the *E. coli* transcriptome.

#### Analysis of essential vs non-essential genes

We first examined the relationship between GCI and individual genes by labelling each gene as either “essential” or “non-essential.” Genes were classified based on their designation in the PEC (Profiling of *E. coli* Chromosome) database (Hashimoto et al. 2005; Yamazaki et al. 2008). We designated genes as essential if they were present in the reference set (*n* = 285), while all other genes were designated as non-essential (*n* = 3625). We tested for a significant difference in mean GCI between these two gene sets using a Student’s t-test.

We checked for robustness of our result with an alternative set of essential genes, defined by the Keio collection (Baba et al. 2006). We extracted all essential genes listed in their Supplementary Table 7 to generate an independent set of essential and non-essential genes (*n* = 255 and *n* = 3655, respectively). We again used a t-test to test for a significant difference in mean GCI between these two sets of genes.

#### Analysis of core vs accessory genes

As an alternative to the distinction of essential and non-essential genes, we also classified all genes in the *E. coli* genome as either “core” or “accessory.” We developed this classification via a Prokka-Roary pipeline, where we identified genes that are shared across the genomes of 60 *E. coli* strains (Seemann 2014; Page et al. 2015; Maddamsetti et al. 2017). This resulted in lists of *n* = 1810 core and *n* = 2100 accessory genes. These numbers were comparable to previously reported results (Abram et al. 2021; Maddamsetti et al. 2017; Sims and Kim 2011). As before, we tested for a significant difference in mean GCI between these two gene sets using a t-test.

#### GO term enrichment analysis

We performed GO term enrichment analysis for the sets of genes with either GCI *<* 0 or GCI *>* 0. For each set, we used the AnnotationDbi package to convert gene names to Entrez IDs, then used the clusterProfiler package to create gene sets of Gene Ontology (GO) terms associated with the gene IDs, and completed an Over-Representation Analysis to find terms enriched in the set (Pagès et al. 2022; Yu et al. 2012; Wu et al. 2021; Ashburner et al. 2000; Gene Ontology Consortium 2021). We only retained terms with significant enrichment (*p* ≤0.04 and *q* ≤ 0.05) and corrected for multiple testing with the Benjamini-Hochberg procedure (Benjamini and Hochberg 1995). Using the output of clusterProfiler’s enrichment analysis, we calculated the odds ratio (OR) for all significant GO terms. For both gene sets, we ranked terms by ORs and reported the terms with the 15 largest ratios.

## Code and data availability

Data analysis, processing, and visualization was done using a combination of python and R scripts (Van Rossum and Drake 2009; R Core Team 2021). These scripts and their associate input and output data are available at: https://github.com/mmjohn/Growth-expression-translation.

## Acknowledgements

This work was supported by National Institutes of Health grant R01 GM088344 to C.O.W. and National Institutes of Health grant F32 GM130113 to A.J.H. C.O.W. also received support from the Jane and Roland Blumberg Centennial Professorship in Molecular Evolution and the Dwight W. and Blanche Faye Reeder Centennial Fellowship in Systematic and Evolutionary Biology at The University of Texas at Austin.

