## Supplementary Figures for "Growth-dependent gene expression variation influences the strength of codon usage biases"

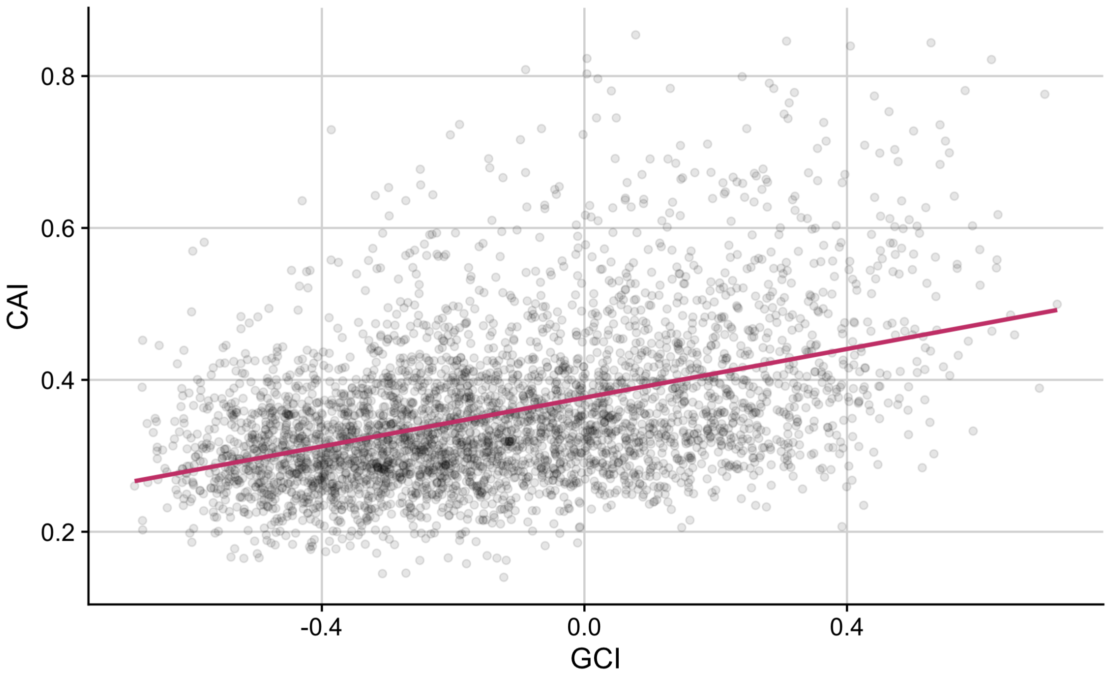


Supplementary Figure S1: **GCI has a significant linear relationship with CAI.** For each gene in the full *E. coli* data set, CAI values are shown against GCI values. The performance of this model relative to models incorporating mean expression data is shown in Figure 5 in the main text.

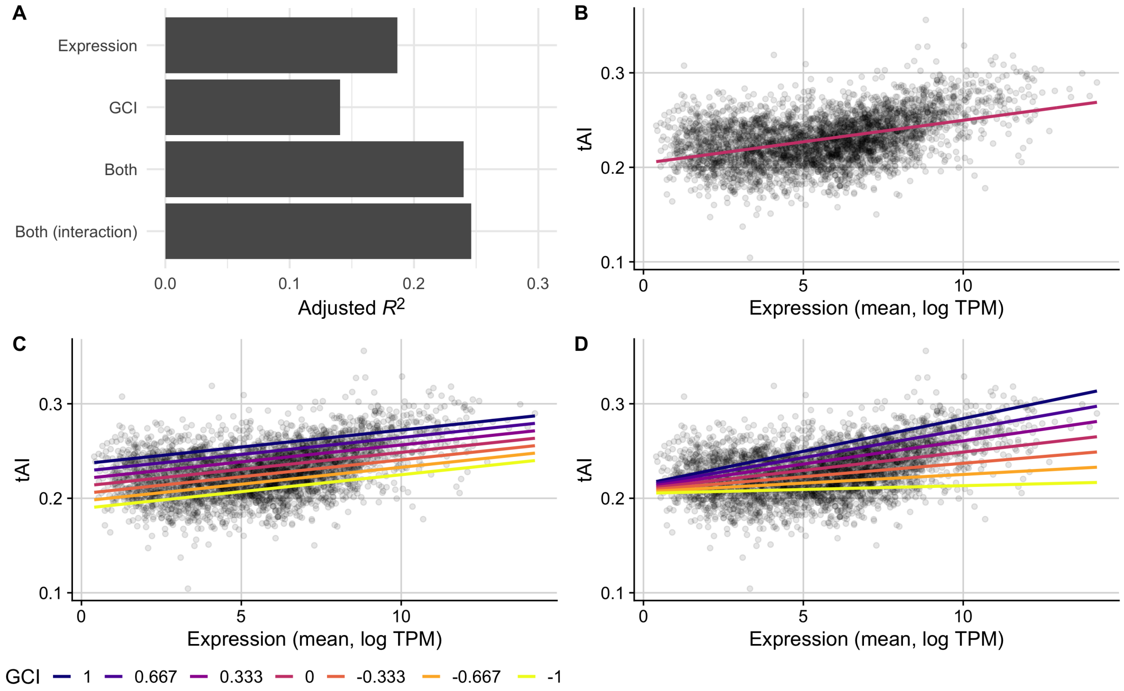


Supplementary Figure S2: **tAI is partially predicted by GCI in the full *E. coli* RNA data set.** A comparison of the predictive ability (measured as $R^2$ after adjustment) of linear models that use either: 1) mean expression values, 2) GCI values, 3) both expression and GCI values, or 4) both values with an interaction term, to predict tAI. (B, C, D) tAI against mean expression for the top 3 performing models with observed values for each gene shown as points and model predictions as lines. The fit of model 1, which predicts tAI using only mean gene expression values, is shown with one line (B), while models 3 and 4 are shown with several lines colored by potential fixed GCI values (C and D, respectively).

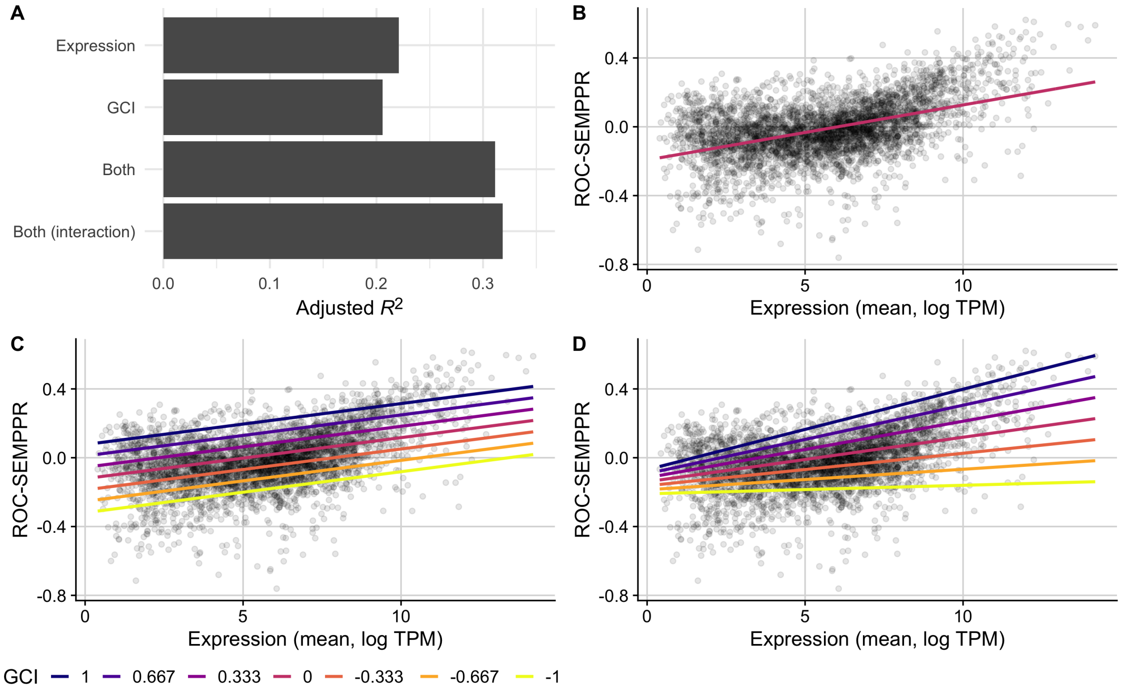


Supplementary Figure S3: **ROC SEMPPR is partially predicted by GCI in the full *E. coli* RNA data set.** A comparison of the predictive ability (measured as $R^2$ after adjustment) of linear models that use either: 1) mean expression values, 2) GCI values, 3) both expression and GCI values, or 4) both values with an interaction term, to predict ROC SEMPPR. (B, C, D) ROC SEMPPR against mean expression for the top 3 performing models with observed values for each gene shown as points and model predictions as lines. The fit of model 1, which predicts ROC SEMPPR using only mean gene expression values, is shown with one line (B), while models 3 and 4 are shown with several lines colored by potential fixed GCI values (C and D, respectively).

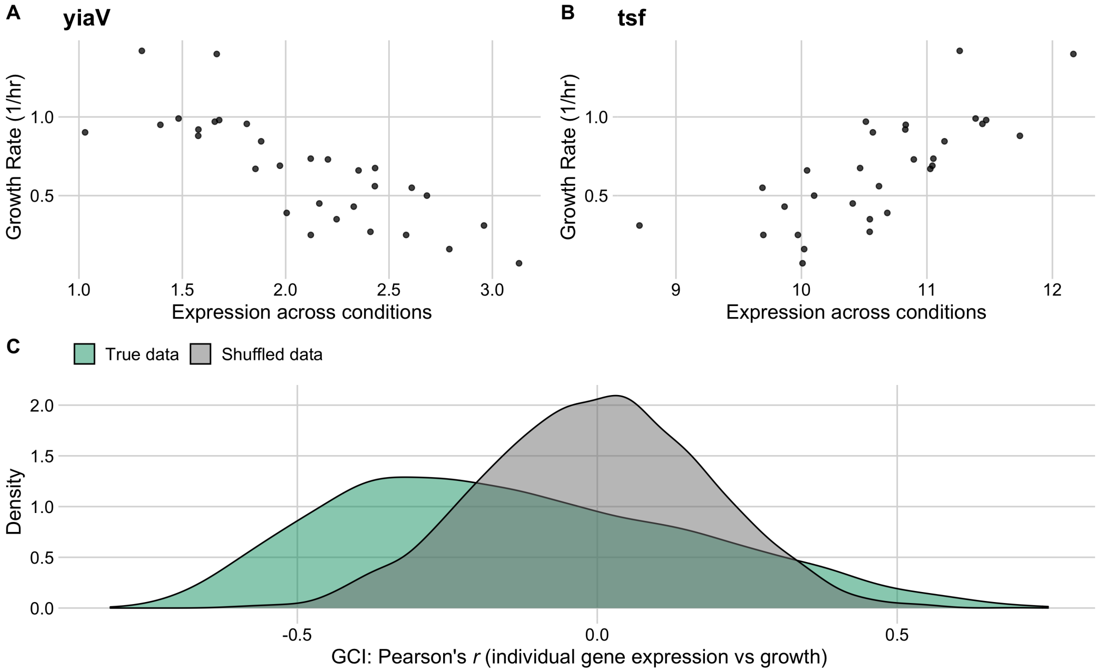


Supplementary Figure S4: **Individual gene expression across conditions variably correlates with growth rate in the sparse *E. coli* RNA data set.** The top row shows the two genes with the most negative (yiaV, A) and most positive (tsf, B) correlation between growth rate and expression across all conditions. (C) The distribution of correlation coefficients for all genes (GCI, shown in green) against a reference data set with permuted expression and growth data (grey).

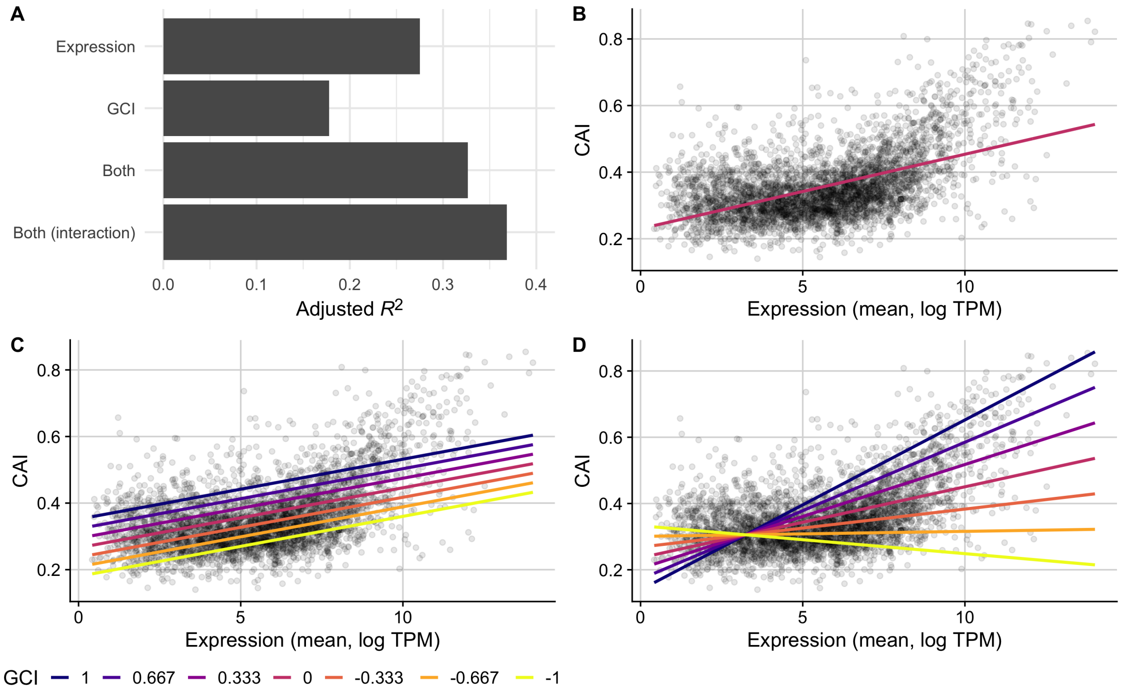


Supplementary Figure S5: **CAI is partially predicted by GCI in the sparse *E. coli* RNA data set.** (A) A comparison of the predictive ability (measured as $R^2$ after adjustment) of linear models that use either: 1) mean expression values, 2) GCI values, 3) both expression and GCI values, or 4) both values with an interaction term, to predict CAI. (B, C, D) CAI against mean expression for the top 3 performing models with observed values for each gene shown as points and model predictions as lines. The fit of model 1, which predicts CAI using only mean gene expression values, is shown with one line (B), while models 3 and 4 are shown with several lines colored by potential fixed GCI values (C and D, respectively). VIF between mean expression and GCI is calculated to be 1.20.

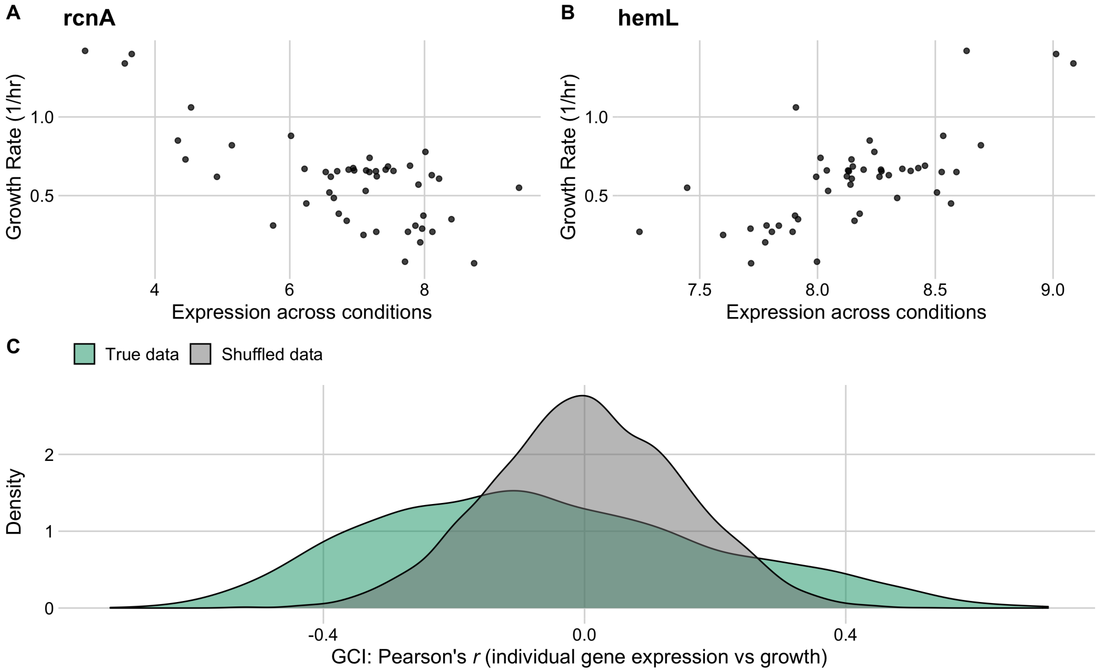


Supplementary Figure S6: **Individual gene expression across conditions variably correlates with growth rate in the neutral *E. coli* RNA data set that excludes ALEs.** The top row shows the two genes with the most negative (rcnA, A) and most positive (hemL, B) correlation between growth rate and expression across all conditions. (C) The distribution of correlation coefficients for all genes (GCI, shown in green) against a reference data set with permuted expression and growth data (grey).

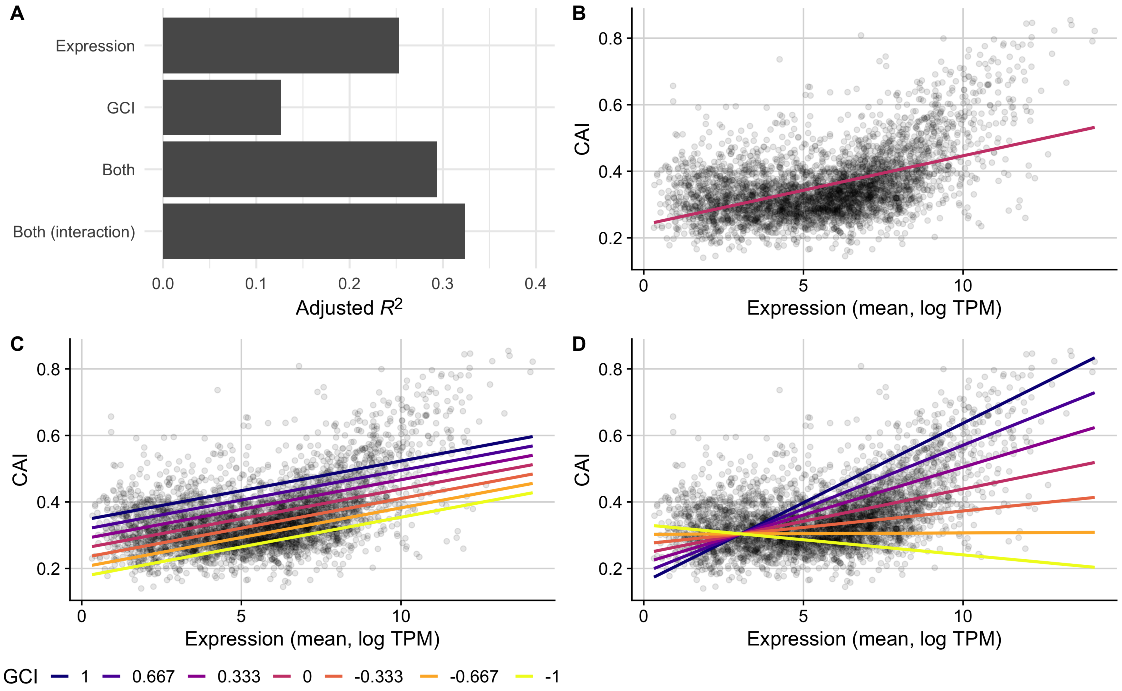


Supplementary Figure S7: **CAI is partially predicted by GCI in the neutral *E. coli* RNA data set that excludes ALEs.** (A) A comparison of the predictive ability (measured as $R^2$ after adjustment) of linear models that use either: 1) mean expression values, 2) GCI values, 3) both expression and GCI values, or 4) both values with an interaction term, to predict CAI. (B, C, D) CAI against mean expression for the top 3 performing models with observed values for each gene shown as points and model predictions as lines. The fit of model 1, which predicts CAI using only mean gene expression values, is shown with one line (B), while models 3 and 4 are shown with several lines colored by potential fixed GCI values (C and D, respectively). VIF between mean expression and GCI is calculated to be 1.19.

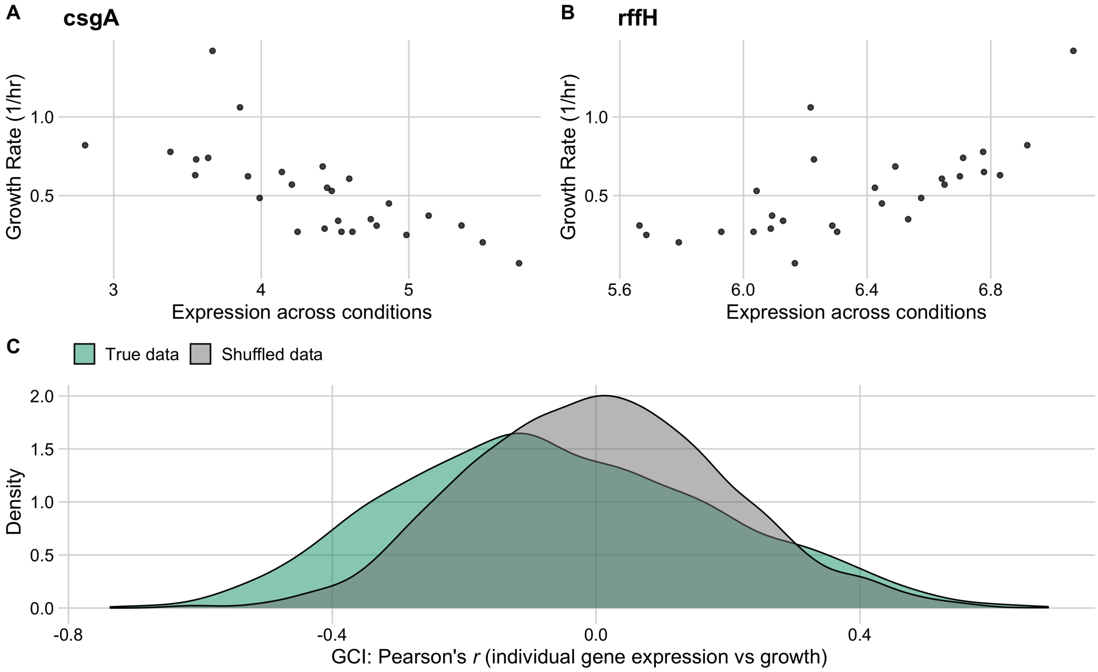


Supplementary Figure S8: **Individual gene expression across conditions variably correlates with growth rate in the neutral *E. coli* RNA data set that excludes ALEs, mutants, and knock-outs.** The top row shows the two genes with the most negative (csgA, A) and most positive (rffH, B) correlation between growth rate and expression across all conditions. (C) The distribution of correlation coefficients for all genes (GCI, shown in green) against a reference data set with permuted expression and growth data (grey).

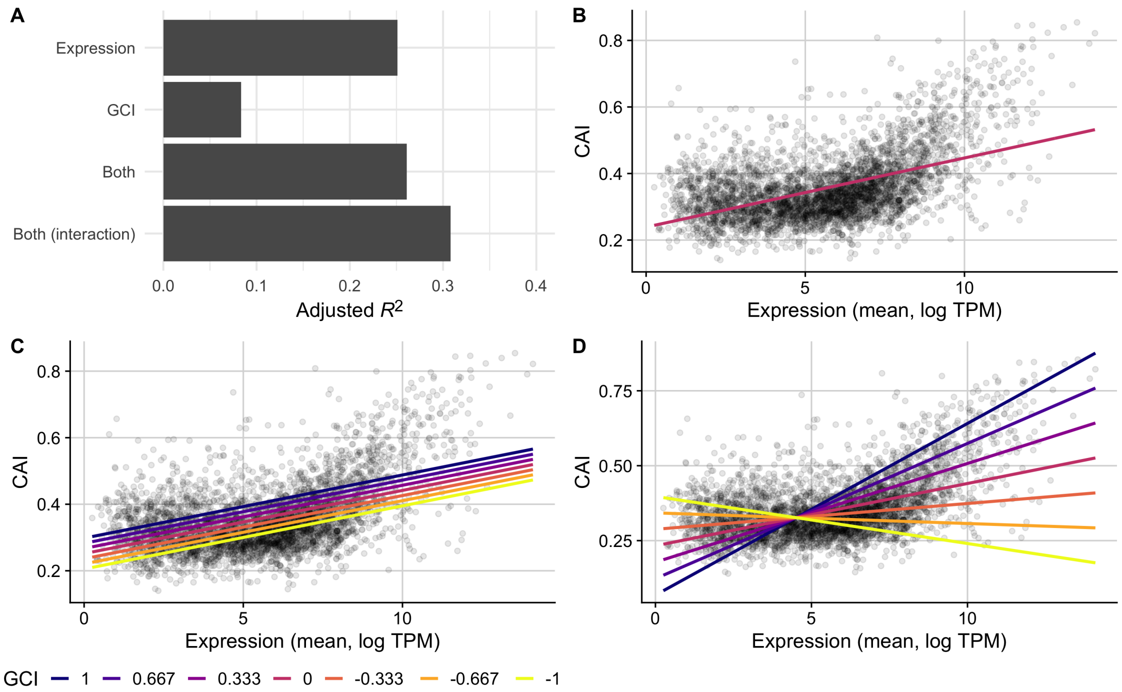


Supplementary Figure S9: **CAI is partially predicted by GCI in the neutral *E. coli* RNA data set that excludes ALEs, knock-outs, and mutants.** (A) A comparison of the predictive ability (measured as $R^2$ after adjustment) of linear models that use either: 1) mean expression values, 2) GCI values, 3) both expression and GCI values, or 4) both values with an interaction term, to predict CAI. (B, C, D) CAI against mean expression for the top 3 performing models with observed values for each gene shown as points and model predictions as lines. The fit of model 1, which predicts CAI using only mean gene expression values, is shown with one line (B), while models 3 and 4 are shown with several lines colored by potential fixed GCI values (C and D, respectively). VIF between mean expression and GCI is calculated to be 1.18.

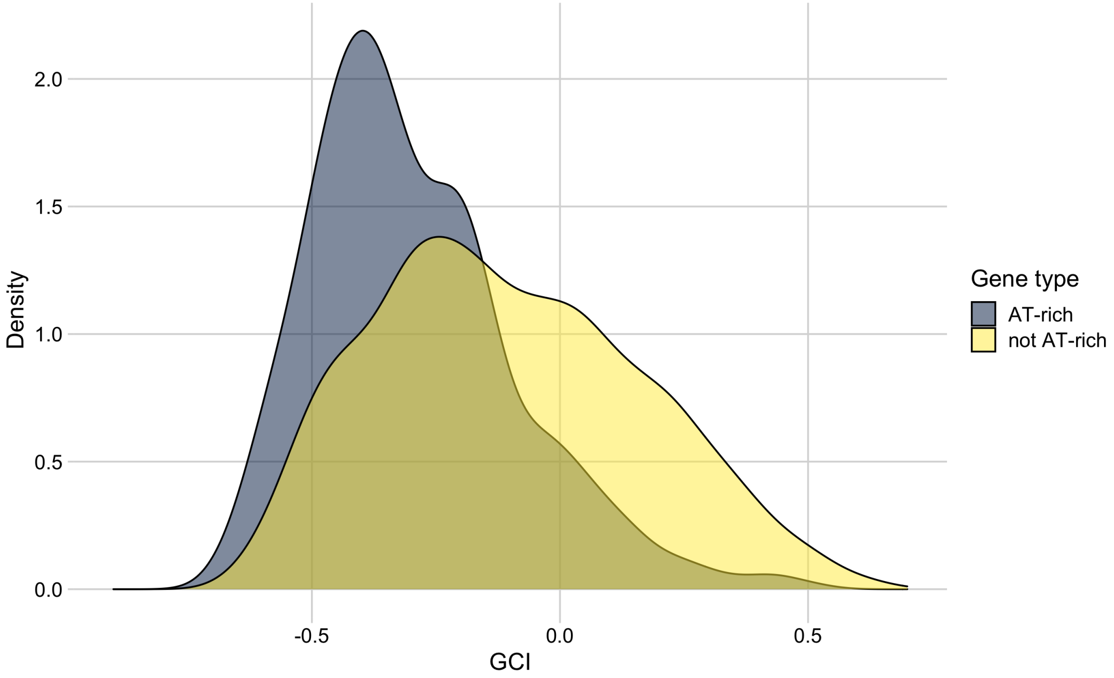


Supplementary Figure S10: **AT-rich genes are skewed towards negative GCI values in the full *E. coli* RNA data set.** The distribution of GCI values for group 3, AT-rich genes from dos Reis et al. (2003) (shown in blue) and for all other genes (yellow). The mean GCI values for the two distributions are $-0.30$ and $-0.11$, respectively, and they are significantly different (t-test, $p < 10^{-10}$).

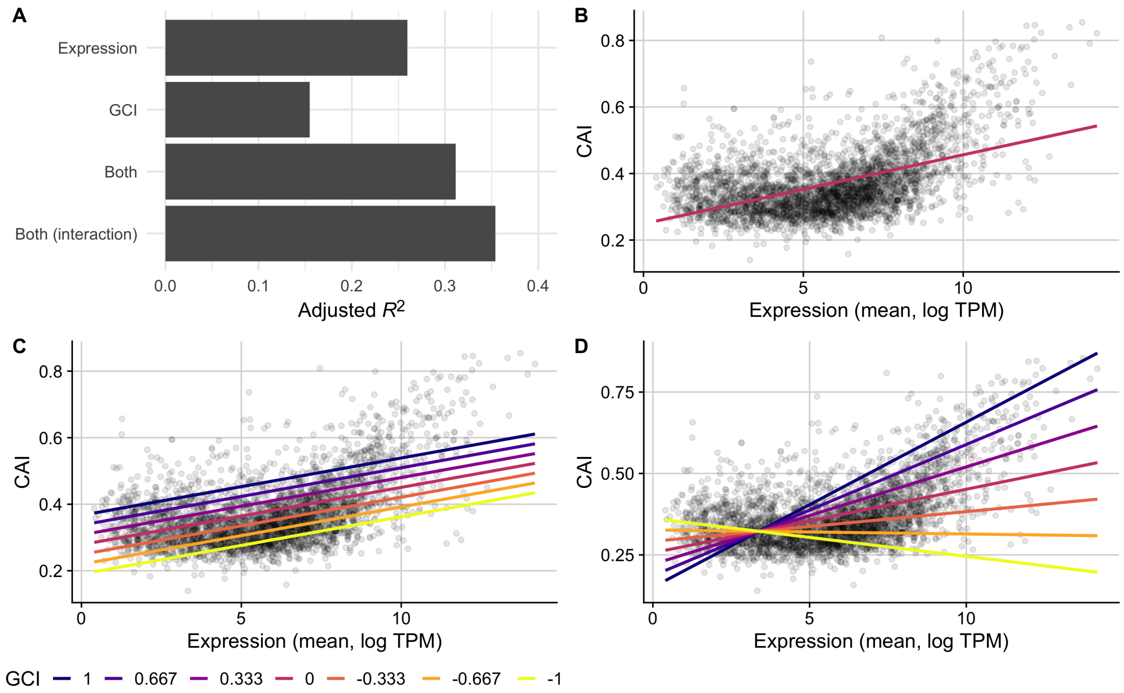


Supplementary Figure S11: **CAI is partially predicted by GCI in the *E. coli* RNA data set that excludes AT-rich genes from dos Reis et al. (2003).** (A) A comparison of the predictive ability (measured as $R^2$ after adjustment) of linear models that use either: 1) mean expression values, 2) GCI values, 3) both expression and GCI values, or 4) both values with an interaction term, to predict CAI. (B, C, D) CAI against mean expression for the top 3 performing models with observed values for each gene shown as points and model predictions as lines. The fit of model 1, which predicts CAI using only mean gene expression values, is shown with one line (B), while models 3 and 4 are shown with several lines colored by potential fixed GCI values (C and D, respectively). VIF between mean expression and GCI is calculated to be 1.15.

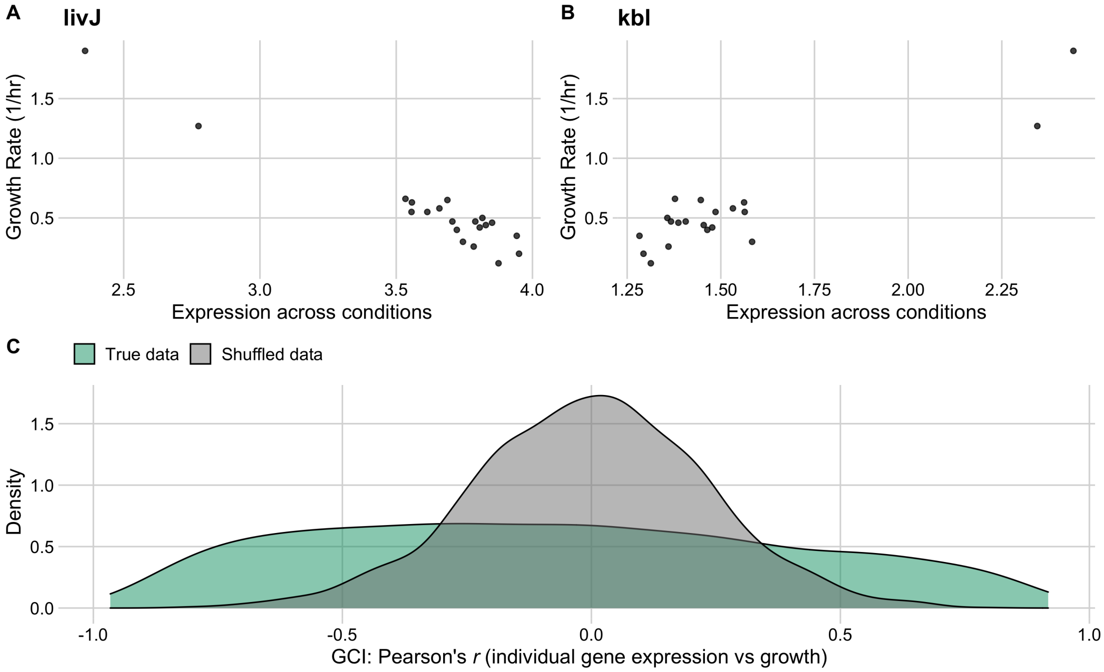


Supplementary Figure S12: **Individual gene expression across conditions variably correlates with growth rate in the *E. coli* protein data set.** The top row shows the two genes with the most negative (livJ, A) and most positive (kbl, B) correlation between growth rate and expression across all conditions. (C) The distribution of correlation coefficients for all genes (GCI, shown in green) against a reference data set with permuted expression and growth data (grey).

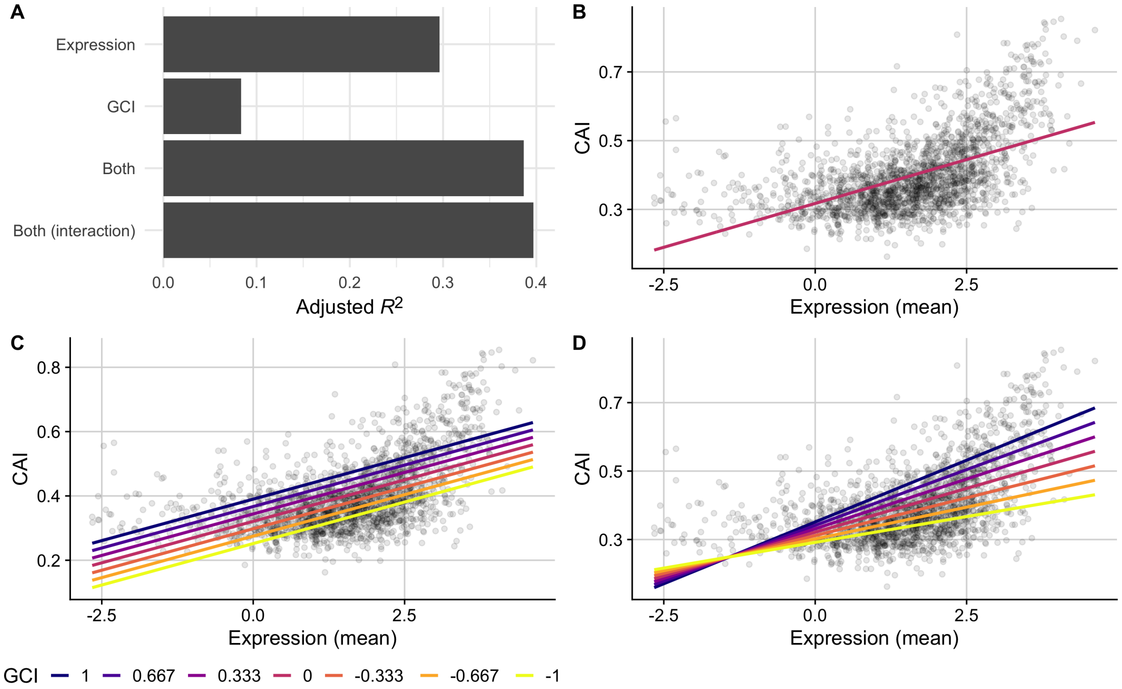


Supplementary Figure S13: **CAI is partially predicted by GCI in the *E. coli* protein data set.** A comparison of the predictive ability (measured as $R^2$ after adjustment) of linear models that use either: 1) mean expression values, 2) GCI values, 3) both expression and GCI values, or 4) both values with an interaction term, to predict CAI. (B, C, D) CAI against mean expression for the top 3 performing models with observed values for each gene shown as points and model predictions as lines. The fit of model 1, which predicts CAI using only mean gene expression values, is shown with one line (B), while models 3 and 4 are shown with several lines colored by potential fixed GCI values (C and D, respectively). VIF between mean expression and GCI is calculated to be 1.00.

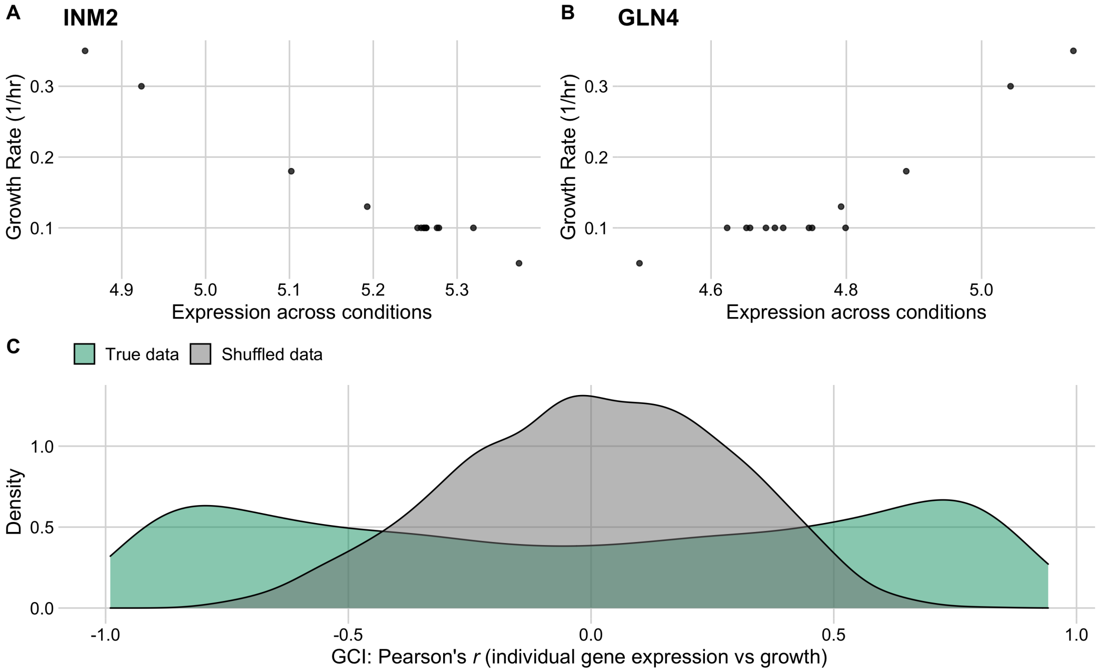


Supplementary Figure S14: **Individual gene expression across conditions variably correlates with growth rate in the *S. cerevisiae* RNA data set.** The top row shows the two genes with the most negative (INM2, A) and most positive (GLN4, B) correlation between growth rate and expression across all conditions. (C) The distribution of correlation coefficients for all genes (GCI, shown in green) against a reference data set with permuted expression and growth data (grey).

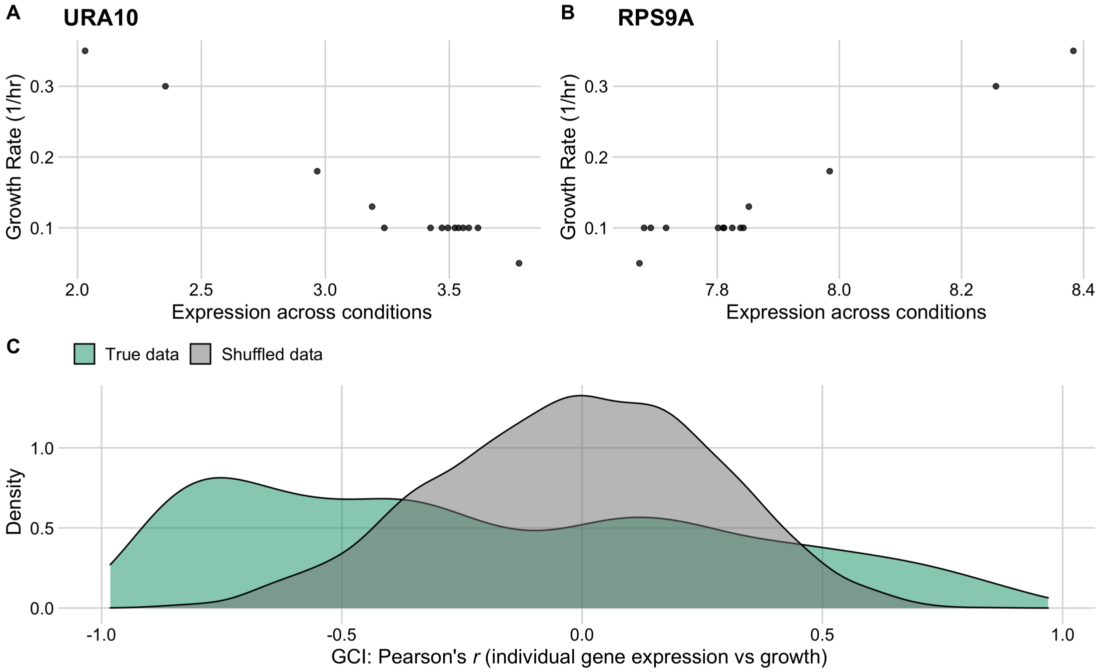


Supplementary Figure S15: **Individual gene expression across conditions variably correlates with growth rate in the *S. cerevisiae* protein data set.** The top row shows the two genes with the most negative (URA10, A) and most positive (RPS9A, B) correlation between growth rate and expression across all conditions. (C) The distribution of correlation coefficients for all genes (GCI, shown in green) against a reference data set with permutated expression and growth data (grey).

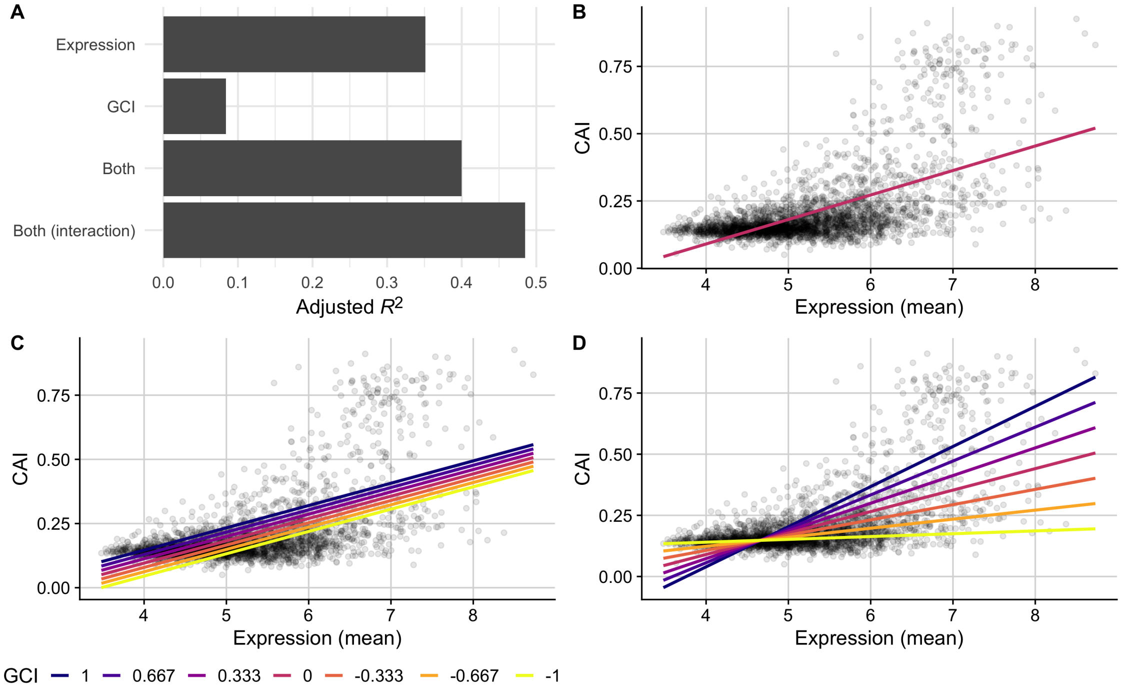


Supplementary Figure S16: **CAI is partially predicted by GCI in the *S. cerevisiae* RNA data set.** A comparison of the predictive ability (measured as $R^2$ after adjustment) of linear models that use either: 1) mean expression values, 2) GCI values, 3) both expression and GCI values, or 4) both values with an interaction term, to predict CAI. (B, C, D) CAI against mean expression for the top 3 performing models with observed values for each gene shown as points and model predictions as lines. The fit of model 1, which predicts CAI using only mean gene expression values, is shown with one line (B), while models 3 and 4 are shown with several lines colored by potential fixed GCI values (C and D, respectively). VIF between mean expression and GCI is calculated to be 1.01.

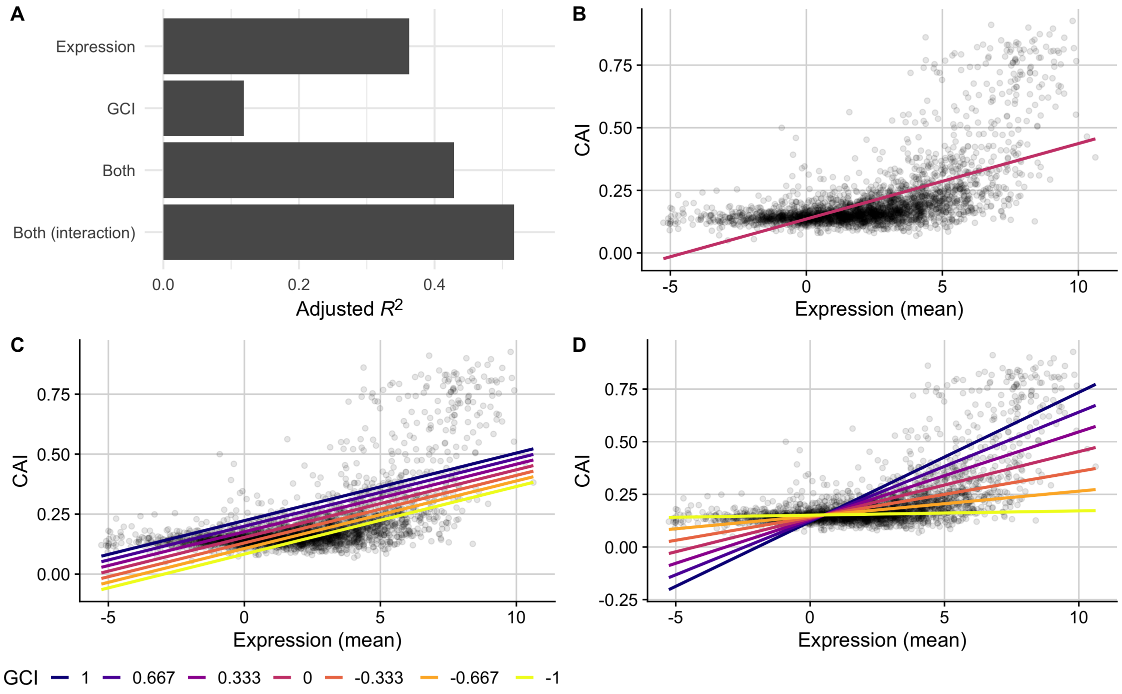


Supplementary Figure S17: **CAI is partially predicted by GCI in the *S. cerevisiae* protein data set.** A comparison of the predictive ability (measured as $R^2$ after adjustment) of linear models that use either: 1) mean expression values, 2) GCI values, 3) both expression and GCI values, or 4) both values with an interaction term, to predict CAI. (B, C, D) CAI against mean expression for the top 3 performing models with observed values for each gene shown as points and model predictions as lines. The fit of model 1, which predicts CAI using only mean gene expression values, is shown with one line (B), while models 3 and 4 are shown with several lines colored by potential fixed GCI values (C and D, respectively). VIF between mean expression and GCI is calculated to be 1.02.
